# RECIFS: a centralized geo-environmental database for coral reef research and conservation

**DOI:** 10.1101/2022.10.20.513055

**Authors:** Oliver Selmoni, Gaël Lecellier, Véronique Berteaux-Lecellier, Stéphane Joost

## Abstract

Host to intricated networks of marine species, coral reefs are among the most biologically diverse ecosystems on Earth. Over the past decades, major degradations of coral reefs have been observed worldwide, which is largely attributed to the effects of climate change and local stressors related to human activities. Now more than ever, characterizing how the environment shapes the dynamics of the reef ecosystem is key to (1) uncovering the environmental drivers of reef degradation, and (2) enforcing efficient conservation strategies in response. To achieve these objectives, it is pivotal that environmental data characterizing such ecosystem dynamics, which occur across specific spatial and temporal scales, are easily accessible to coral reef researchers and conservation stakeholders alike.

Here we present the Reef Environment Centralized Information System (RECIFS), an online repository of datasets describing reef environments worldwide over the past few decades.

The data served through RECIFS originate from remote sensed datasets available in the public domain, and characterize various facets of the reef environment, including water chemistry and physics (e.g. temperature, pH, chlorophyll concentration), as well as anthropogenic local pressures (e.g. boat detection, distance from agricultural or urban areas). The datasets from RECIFS can be accessed at different spatial and temporal resolutions and are delivered through an intuitive web-application featuring an interactive map requiring no prior knowledge working with remote sensing or geographic information systems. The RECIFS web-application is available in complete open access at https://recifs.epfl.ch.

We describe two case studies showing possible implementations of RECIFS to 1) characterize coral diversity in the Caribbean and 2) investigate local adaptation of a reef fish population in Northwest Australia.

## Introduction

Host of up to a quarter of marine species, coral reefs are among the most productive and biologically diverse ecosystems on Earth (Bouchet, 2006; Knowlton et al., 2010; Reaka-Kudla, 1997). These biodiversity hotspots are currently facing critical threats imposed by both climate change and human-induced local disturbances (Hughes et al., 2017). Indeed, a 14% loss of hard coral cover worldwide was recently estimated for the 2009-2018 decade, where the decline was mainly attributed to anomalous heat waves causing coral bleaching (Souter et al., 2021). Additionally, the resilience of corals to thermal stress can be hampered by local water conditions, such as turbidity levels or nutrient loads (MacNeil et al., 2019). As hard corals shape the physical reef habitat, their loss can lead declines in reef fish abundance (G. P. Jones et al., 2004), and such declines can be exacerbated by local stressors such as overfishing (Hughes et al., 2017) and pollution (e.g. agricultural or industrial runoff; Wenger et al., 2015). Overall, the deterioration of coral reefs impairs the associated ecosystem services, which are key for the well-being of human communities in the tropics (Eddy et al., 2021).

To identify the drivers and quantify rates coral reef deterioration, researchers can investigate the association between remote sensed environmental data, and *in-situ* measurements replicated at multiple locations across reef systems. These *in-situ* measurements can include various ecological surveys, including surveys of coral abundance (e.g., Sully et al., 2022), coral diversity (e.g., Kusumoto et al., 2020), coral bleaching severity (e.g., McClanahan et al., 2020), fish biomass (e.g., Cinner et al., 2016), or the presence/absence of a given reef species (e.g., Förderer et al., 2018; Ottimofiore et al., 2017; Principe et al., 2021). *In-situ* measurements also include molecular data from populations that represent genetic diversity, which can be used in genotype-environment association (GEA) studies to uncover genetic variants potentially underpinning local adaptation processes (Fuller et al., 2020; Lundgren et al., 2013; Selmoni, Lecellier, Magalon, et al., 2020; Selmoni, Rochat, et al., 2020; Sherman et al., 2020; Thomas et al., 2017).

These association studies can expose the environmental drivers of species abundance, diversity and adaptive potential, which are all pivotal elements to organize effective reef conservation strategies (Foo & Asner, 2019, 2021; Hedley et al., 2016; Lecours et al., 2021; Murray et al., 2018). For example, high taxonomic diversity is among the main criteria used to prioritize reefs when establishing marine protected areas (MPAs; Mascia et al., 2017). Additionally, MPAs could be enforced at reefs with a thermal history associated with heat-adaptation in corals (Wilson et al., 2020). The optimal location of these MPAs can then be decided based on regional surface water circulation, such that larval dispersal of thermally adapted corals reaches the largest number of reefs downstream of protected populations (Matz et al., 2020; Selmoni, Lecellier, Vigliola, et al., 2020; Selmoni, Rochat, et al., 2020; van Woesik et al., 2022).

For conservation strategies to make the most of these environmental drivers, the underlying environmental data must be easily accessible for both the researchers running the association analyses and the decision makers enforcing reef conservation strategies (Hedley et al., 2016; Lecours et al., 2021). There are already online repositories that provide synthesized variables of marine environments worldwide, such as Bio-ORACLE (Assis et al., 2018; Tyberghein et al., 2012) and MARSPEC (Sbrocco & Barber, 2013). The synthesized variables are derived from remotely sensed data obtained from public initiatives, such as the World Ocean Atlas (Boyer et al., 2018) and the Copernicus Marine Service (EU Copernicus Marine Service, 2022), and describe long-lasting trends in global oceanic conditions. These synthesized datasets are predominantly produced (and widely used) for spatial distribution modelling of marine organisms (Sbrocco & Barber, 2013; Tyberghein et al., 2012).

However, these repositories are not coral reef-centered, and therefore do not give access to environmental variables that are key to understand the dynamics of the coral reef ecosystem, such as coral heat stress indices (e.g., those from the Coral Reef Watch; Liu et al., 2014) or indices of local fishing activities or of human land use along the coastline. In addition, these repositories offer limited possibilities for customizing the spatial and temporal extents of environmental data, where such customization is key when analyzing the relationship with *in-situ* measurements (Leempoel et al., 2015; Murray et al., 2018). For example, a certain *in-situ* measurement (e.g., the abundance of a given species; the presence of an adaptive genetic trait) could be driven by an environmental variation during a specific season (e.g., thermal stress during the hot season), or by pollution-related human activities within a certain distance from the coastline (e.g., agricultural areas in a 50 km radius). As the sub-setting and processing of environmental data is generally performed using dedicated software (e.g., R-packages, Python), this data processing generally requires competences with geographic information systems (GIS) and scientific programming. The lack of an interactive and easy-to-use user interface hampers the integration of these data by non-GIS experts, as may be the case for coral reef conservation practitioners (Guisan et al., 2013; Selmoni, Lecellier, Ainley, et al., 2020).

Here, we present the Reef Environment Centralized Information System (RECIFS), an online repository of datasets describing the reef environment worldwide over the past decades. The datasets provided through RECIFS originate from the public domain, and characterize various facets of the reef environment including water chemistry (e.g., chlorophyll concentration, salinity, etc.) and physics (e.g., temperature, heat, velocity, etc.), as well as anthropogenic impacts (e.g., population density, land use of agricultural and urban areas) along the neighboring coastlines. The datasets from RECIFS can be accessed at different spatial resolutions and are sub-settable along the temporal dimension (*i.e.*, select specific seasons and/or specific years). The data are delivered through an intuitive a free-to-use web-GIS application, available at https://recifs.epfl.ch. We hereunder present the functionalities of this tool, and describe two cases studies showing possible implementations: 1) the characterization of coral diversity in the Caribbean, and 2) the study of local adaption of a reef fish population along the northwestern coast of Australia.

## Methods

### Data structure: reef environment

The RECIFS repository is based on three types of datasets describing the reef environment retrospectively at different temporal resolutions: monthly, annual and non-temporal (i.e., variables without a temporal resolution; Table 1). These datasets were accessed at a global level as raw data, and were then processed and synthesized as outlined here below (and as summarized in Figure 1).

**Table 1.** Datasets included in RECIFS. For each dataset, the table shows the name and description of the variable of interest, the temporal window covered by the dataset (with the temporal resolution in parenthesis), the spatial resolution of the dataset, the source repository of the dataset (with the abbreviations explicited in the bottom) and the identifier of the original product from the source repository.

| Name | Description | Time window (resolution) | Spatial res. | Source | Original product id |
| --- | --- | --- | --- | --- | --- |
| Chlorophyll concentration [mg m<sup>-3</sup>] | Monthly averages of mass concentration of chlorophyll-a in seawater. | 1997-2021 (monthly) | 4 km | CMEMS | OCEANCOLOUR_GLO_CHL_L4_REP_OBSERVATIONS_009_082 |
| Degree heating week [°C-week] | Monthly maxima of degree heating week (DHW). DHW is calculated as the accumulation of thermal stress (i.e. temperature >1°C above the monthly maximal mean temperature) over the previous 12 weeks. | 1985-2021 (monthly) | 5 km | NOAA CRW | ct5km_dhw-max_v3.1 |
| Iron concentration [mmol m<sup>-3</sup>] | Monthly averages of mole concentration of dissolved iron in sea water. | 1993-2019 (monthly) | 0.25 deg | CMEMS | GLOBAL_MULTIYEAR_BGC_001_029 |
| O2 concentration [mmol m<sup>-3</sup>] | Monthly averages of mole concentration of dissolved molecular oxygen in sea water. | 1993-2019 (monthly) | 0.25 deg | CMEMS | GLOBAL_MULTIYEAR_BGC_001_029 |
| Sea water pH | Monthly averages of sea water pH reported on total scale. | 1993-2019 (monthly) | 0.25 deg | CMEMS | GLOBAL_MULTIYEAR_BGC_001_029 |
| Nitrate concentration [mmol m<sup>-3</sup>] | Monthly averages of mole concentration of nitrate in sea water. | 1993-2019 (monthly) | 0.25 deg | CMEMS | GLOBAL_MULTIYEAR_BGC_001_029 |
| Phosphate concentration [mmol m<sup>-3</sup>] | Monthly averages of mole concentration of phosphate in sea water. | 1993-2019 (monthly) | 0.25 deg | CMEMS | GLOBAL_MULTIYEAR_BGC_001_029 |
| Sea water velocity [m s<sup>-1</sup>] | Monthly averages of sea water surface velocity, computed as the Euclidean norm of eastward and northward velocity. | 1993-2019 (monthly) | 0.083 deg | CMEMS | GLOBAL_REANALYSIS_PHY_001_030 |
| Suspended matter concentration [g m<sup>-3</sup>] | Monthly averages of mass concentration of suspended matter in sea water. | 1997-2021 (monthly) | 4 km | CMEMS | OCEANCOLOUR_GLO_OPTICS_L4_REP_OBSERVATIONS_009_081 |
| Sea surface salinity [1e-3] | Monthly averages of sea surface salinity. | 1993-2019 (monthly) | 0.083 deg | CMEMS | GLOBAL_REANALYSIS_PHY_001_030 |
| Sea surface temperature [°C] | Monthly average of sea surface temperature. | 1985-2021 (monthly) | 5 km | NOAA CRW | ct5km_sst-mean_v3.1 |
| Density of built-up [%] | Percentage of ground cover of built-up areas per pixel | 2015-2019 (yearly) | 100 m | CGLS | LandCover100m:collection_3:epoch2019 |
| Density of cropland [%] | Percentage of ground cover of cropland per pixel | 2015-2019 (yearly) | 100 m | CGLS | LandCover100m:collection_3:epoch2019 |
| Boat detection [%] | Percentage of boat detections per satellite overpass coverage | 2017-2020 (yearly) | 0.04 deg | EOG | VBD_npp_global-saa_pc_v23 |
| Human population density [hab km<sup>-2</sup>] | Human population density | 2000-2020 (yearly - by 5 years) | 0.04 deg | SEDAC | gpw-v4- rev11 |
| Depth [m] | Depth | - | 0.017 deg | GMRT | GMRTv3_9 |
| Surface land | Binary map, indicating whether pixel is on land or sea. | - | 0.017 deg | GMRT | GMRTv3_9 |
| **Data sources** |  |  |  |  |  |
| Source id | Source name | Reference |  |  |  |
| CMEMS | Copernicus Marine Service | (EU Copernicus Marine Service, 2022) |  |  |  |
| NOAA CRW | National Oceanic and Atmospheric Administration, Coral Reef Watch | (NOAA Coral Reef Watch, 2022) |  |  |  |
| CGLS | Copernicus Global Land Service | (Copernicus Global Land Service, 2022) |  |  |  |
| EOG | Earth Observation Group - Payne Institute for Public Policy | (Elvidge et al., 2015, 2018; Hsu et al., 2019) |  |  |  |
| SEDAC | Socioeconomic Data and Applications Center, Columbia University | (Center for International Earth Science Information Network, 2022) |  |  |  |
| GMRT | Global Multi-Resolution Topography Data Synthesis | (Ryan et al., 2009) |  |  |  |

**Figure 1.**
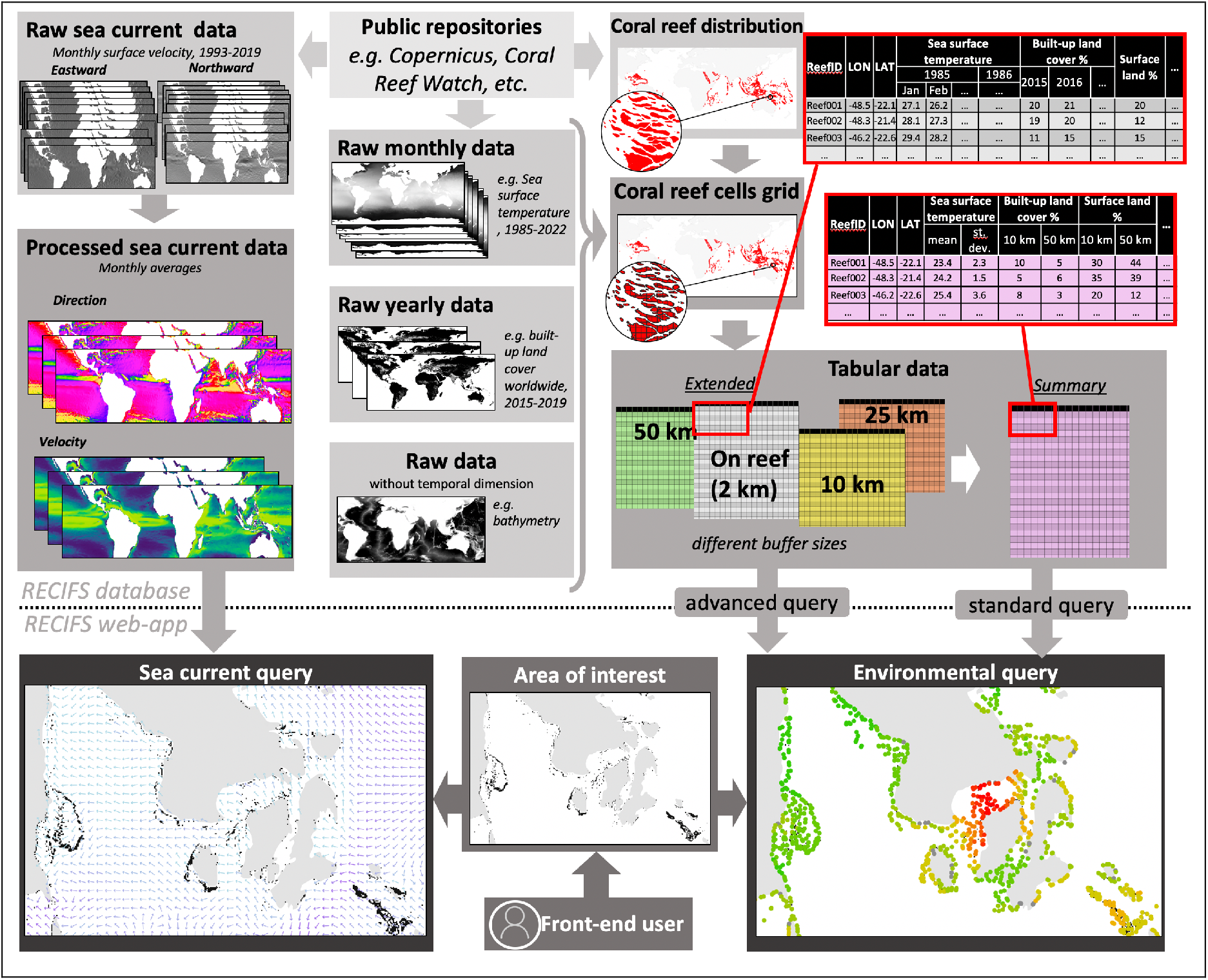
Data workflow. Raw environmental data used in RECIFS comes from publicly available repositories, and have distinct temporal resolutions (monthly, annual or no resolution). Shapes of coral reefs worldwide were summarized into a regular grid (reef cell size: 5-by-5 km), and raw environmental values were extracted for each reef cell. These extractions were performed using four different buffer sizes (2.5 km – on reef, 10 km, 25 km or 50 km) around the centers of reef cells. The result is a set of tables (“Extended tables”) describing variation across reef cells for each environmental variable, at different buffer sizes and through different time periods (months or years). These tables are synthesized into a “Summary table”, reporting overall temporal average and standard deviation of each environmental variable, at every reef cell. RECIFS also includes data on sea surface current velocity and direction, obtained from processed raw data describing eastward and northward surface water velocity. These processed data are accessible through the RECIFS web-application: the user selects an area of interest through an interactive map and runs either an environmental query to visualize environmental variation (standard query to access the Summary table, advanced query to access the Extended tables), or a sea current query to visualize sea currents.

Raw datasets with a monthly temporal resolution were stacks of raster images, where every image described a monthly statistic (usually the average) of a given oceanic environmental variable globally. RECIFS includes the raw monthly datasets for the following environmental variables: chlorophyll concentration, degree heating week, iron concentration, oxygen concentration, pH, nitrate concentration, phosphate concentration, sea current velocity, suspended matter concentration, salinity, and temperature. These variables represent data at near-surface depth, with spatial resolutions ranging between 5-25 km, and across temporal windows generally covering the past 2-3 decades (see Table 1 for details). These raw monthly datasets are sourced from the repositories from the Coral Reef Watch of the National Oceanic and Atmospheric Administration (NOAA Coral Reef Watch, 2022) and the Copernicus Marine Service (EU Copernicus Marine Service, 2022).

Raw datasets at yearly temporal resolution were also raster stacks, but here each image was a yearly measure of a given variable worldwide. The datasets of this type included in RECIFS were: land cover of built-up and cropland areas (covering the 2015-2019 period, at 100m resolution; Copernicus Global Land Service, 2022), boat detection (2017-2020, at 500m resolution; Elvidge et al., 2015, 2018; Hsu et al., 2019), and human population density (available for the years 2000, 2005, 2010, 2015, 2020, at 5 km resolution, Center for International Earth Science Information Network, 2022).

Two raw datasets without a temporal resolution were included in RECIFS, both derived from the Global Multi-Resolution Topography Data Synthesis, which is a single raster image describing topography (on land) and bathymetry (on sea) worldwide at ∼1 km of spatial resolution (Ryan et al., 2009). The first derived dataset is a depth map where the pixels represent the topography of below sea surface level, and the second is a binary map where pixels were either assigned to land (topography > 0) or to sea (topography < 0).

To process these raw environmental datasets, we extracted values from the raster stacks using custom R scripts (v. 4.1; R Core Team, 2021), featuring the raster (v. 3.5; Hijmans, 2021) and the rgdal (v. 1.5; Bivand et al., 2021) libraries. First, polygons representing coral reefs worldwide were retrieved from the Global Distribution of Coral Reefs dataset (v. 4.1; UNEP-WCMC et al., 2021), and were reported to 5-by-5 km cells. The resulting grid was composed of 61,038 reef cells. For each reef cell, we extracted the environmental values from each raw dataset using a buffer to average over different radiuses (2.5 km, referred to as “on reef” extraction, and 10 km, 25 km and 50 km). The result was a set of tables (called “Extended tables”), with one table per environmental variable at a given buffer size. For each table, the rows represented the worldwide reef cells (such that there were 61,038 rows in each table) and columns represented the temporal dimension (monthly, yearly or a single column for variables without temporal resolution).

Finally, the “Extended tables” were synthesized together into a single table (called the “Summary table”). For the variables at a monthly resolution, each reef cell had overall temporal average and standard deviation “on reef” (i.e., using the 2.5 km buffer) for each environmental variable. For the annual and non-temporal variables, each reef cell had overall averages measured using two of the buffer sizes: 10 and 50 km.

### Data structure: surface currents

The raw environmental dataset used to characterize sea currents was sourced from the Copernicus Marine Service (dataset id: GLOBAL_REANALYSIS_PHY_001_030; EU Copernicus Marine Service, 2022). This dataset is obtained from a stack of raster images displaying global monthly averages of eastward and northward sea surface velocity, from 1993 to 2019. Using custom R-scripts featuring the raster and the circular (v. 0.4; Agostinelli & Lund, 2017) libraries, we combined eastward and northward velocities of each pixel into a vector of sea currents, and then computed (1) sea current direction (the angle of the vector) and (2) sea current velocity (the norm of the vector) for each monthly measure. Finally, we summarized trends of surface circulation by computing raster images representing the overall yearly average and twelve by-month averages for both sea current direction and velocities.

### Web-app and queries

The web-based application of RECIFS is built on a NodeJS server (v. 10.13; OpenJS Foundation). On the front-end, the server features an Openlayers (v 5.3; Open Source Geospatial Foundation) map interface displaying the reef cells grid, while the back-end of the server stores environmental data in a tabular format and the raster images summarizing sea surface currents.

When the front-end user defines an “Area of Interest” through the interactive map, the server queries the RECIFS database through R-scripts (featuring the raster and the sp libraries; v. 1.4; Bivand et al., 2013) processing, sub-setting and finally returning the data stored on the back-end. Three types of query are possible:

1. <u>standard environmental query</u>: the user selects an “Area of Interest”, and the server returns precomputed environmental values from the “Summary table” for the reef cells from this area.
2. <u>advanced environmental query</u>: the user selects an “Area of Interest”, an environmental variable of interest, a spatial buffer size, a temporal window (years and/or months), and a statistic to summarize environmental variation over time (mean, standard deviation, median, minimum or maximum), and the server returns the corresponding values from the “Extended tables” for the reef cells in the area.
3. <u>sea current query</u>: the user selects an “Area of Interest” and specifies a month of interest, and the server returns a visual display of arrows indicating the corresponding strength and direction of sea surface currents.

The results of the queries are displayed on the interactive map. The front-end interface features different functionalities allowing the user to customize map visualization, such as defining the color scale used to display environmental data, setting the transparency of layers or modifying the background layer. The map rendered on the web-interface can be downloaded either in a pdf or a tabular format. Furthermore, the user can define “Points of interest” on the map (e.g., sampling locations, reefs of interest) and download the environmental data for the closest reef cells to such points. All these functionalities are available in complete open access, without any registration required.

## Results & Discussion

### A repository to characterize the reef environment

RECIFS is the first repository that synthesizes data from various sources to describe reef environments globally. In comparison to existing repositories of marine environmental data (Bio-ORACLE, Assis et al., 2018; Tyberghein et al., 2012; MARSPEC, Sbrocco & Barber, 2013), RECIFS has distinctive features to support (1) the study of coral reef ecosystem dynamics, and (2) the organization of reef conservation strategies.

The data provided through RECIFS is centered on the reef environment and characterize key environmental constraints of this ecosystem, including conditions that trigger coral bleaching events and represent coastal pollution-related activities. RECIFS data are available under specific spatial and temporal scales, providing pivotal information for researchers investigating how the reef environment shapes ecosystem dynamics (Melo-Merino et al., 2020; Murray et al., 2018). For instance, modelling the abundance or diversity (e.g., Box 1) of reef taxa with different characteristics (e.g., sessile vs. mobile), or facing different types of environmental constraints (e.g., constant vs. episodic), will require distinct spatial and temporal resolutions of the environmental dataset (Fernandez et al., 2017; Mannocci et al., 2017; Melo-Merino et al., 2020). The same considerations apply to molecular studies on local adaptation (e.g., Box 2), as the genetic makeup of sampled individuals is shaped by ancestors exposed to differing frequencies of environmental stressors (i.e., temporal resolution of variables to be adjusted accordingly), which dispersed over larger or smaller spatial scales (i.e., spatial resolution of variables to be adjusted accordingly; Leempoel et al., 2017; Riginos et al., 2016).

RECIFS data are available in open access through an intuitive interface requiring no prior knowledge on use of geographic information systems. These characteristics make remotely sensed environmental data accessible to non-specialists of the field, such as practitioners of coral reef conservation (Selmoni, Lecellier, Ainley, et al., 2020). Indeed, RECIFS can be a platform to facilitate exchanges between coral reef researchers and conservation practitioners, as it provides a common syntax to describe the environmental factors shaping the dynamics of the reef ecosystem (Guisan et al., 2013). Overall, the goal of RECIFS is to encourage the use of remote sensed data in coral reef conservation strategies worldwide, as advocated by previous review in the domain (Foo & Asner, 2019, 2021; Hedley et al., 2016).

### Perspectives

In the years to come, the development of RECIFS will focus on three domains: data, functionalities and integration. The current data available in the repository will be updated on an annual basis to provide updated records for the different environment variables. Furthermore, new variables that describe environmental factors shaping the dynamics of coral reef species will be added to the database as these variables become available and/or are required; we encourage coral reef researchers and conservation actors to contact us (the authors) to propose environmental descriptors to be added to RECIFS. With the future improvements of remote sensed data and environmental modelling techniques, we also aim to increase the spatial and temporal resolution of the reef cells, so that the environmental characterization can be more pertinent to fine-scale dynamics of the reef ecosystem (Murray et al., 2018). For example, one future goal will be to characterize reef pools located a few hundreds of meters apart, or to characterize daily fluctuations in sea surface temperature (Schoepf et al., 2015; Smith et al., 2007).

Concerning functionalities, the main future goal is to develop an objective quantification of reef connectivity calculated from sea current data. Such quantification could be based on transition matrices and graph theory (such as is implemented in the gdistance R package; van Etten, 2018), and process sea surface current during a particular season to estimate the probability of transition from one reef to another. The combination of such sea current estimates with population genomics data could be used to assess patterns of reef connectivity for a given species. Overall patterns of regional connectivity could then be synthesized in connectivity indices, summarizing how each reef of a given region is interconnected to those located downstream or upstream via oceanic currents (Selmoni, Rochat, et al., 2020). Information on reef connectivity is necessary for undertaking effective conservation efforts, as it informs MPA managers on how to establish networks that best facilitate the larval dispersal to/from key conservation units (e.g. reefs hosting thermally adapted corals; Matz et al., 2020; McCook et al., 2010).

Going forwards, it will also be important to integrate RECIFS with other projects and complementary datasets that characterize alternative facets of reef environments. This could be done, for instance, by linking RECIFS with Reef Cover, which is a classification system based on remote sensing and *in-situ* data that categorizes reefs worldwide by habitat type (e.g. reef slope, reef crest, shallow lagoon, etc.; Kennedy et al., 2021). Importantly, the cross-link between repositories should also be extended to other types of data (i.e., non-environmental) that describe the reef ecosystem (Hedley et al., 2016; Lecours et al., 2021). Such data could be virtually any type of geo-referenced information, for example field surveys on species abundance and diversity, or genomic data characterizing the genetic diversity of a given reef population. The case studies presented here (Box 1 and 2) show how linking different layers of information on the state of the reef can produce valuable insights to orientate conservation strategies. A hub centralizing different data sources on the state of the reef will be of paramount importance in the future, as it will facilitate the establishment of the links between all the different information layers. In the long-term, the goal of RECIFS is to serve as foundation to build such a hub.

#### Box 1. Coral taxonomic diversity in the Caribbean.

The first case study is based on photographic surveys performed across the reefs of the Caribbean by the Catlin Seaview Survey (CSS) project (A; González-Rivero et al., 2014, 2016). The CSS project used machine learning to annotate the presence of 17 hard coral taxa along 183 surveys performed in 12 territories of the region. We used these taxonomic annotations to calculate the Simpson’s diversity index (SDI) for each survey, and then followed the analytical pipeline shown in (B) to investigate the association between spatial variation of SDI and 302 environmental variables obtained from RECIFS.

The 302 variables characterized the reef environment of the region at varying spatial and temporal resolutions. We found that 11 of these variables significantly (p<0.05) explained variation of the SDI (R^2^=0.56). These variables describe environmental conditions that are well known to mediate changes in coral abundance/mortality, such as thermal stress (standard deviation of Degree Heating Week during the wet season), ocean acidification (standard deviation of pH), or run-off from agricultural activities along the coastline (land use of cropland; Cornwall et al., 2021; D’Angelo & Wiedenmann, 2014; De’ath & Fabricius, 2010; R. Jones et al., 2020; McClanahan et al., 2020; Sully et al., 2019). In (C), we show the association between SDI and four of these variables: standard deviation of PH (PH_025_DS_sd), sea surface temperature (SST_050_DS_sd), degree heating week (DHW_010_WS_sd), and density of cropland (CROP_025_me).

Based on the associations described above, the RECIFS data were used to predict SDI for each reef of the Caribbean (D). This spatial prediction highlighted regions expected to host reefs with high coral diversity, for instance in Belize (highlighted by the purple “I” on map); Bonaire, Aruba and Curaçao (“II”); Dominica and Martinique (“III”); and the Andros Island in Bahamas (“IV”). The mean absolute error (MAE) of the prediction was ∼11% of the range of SDI measured in real surveys and did not appear to increase with distance from survey locations. Care must be taken when interpreting this prediction as the survey data was systematically collected at 10 meters of depth, such that these diversity estimations might not be relevant at other depth levels.

Some of the biodiversity patterns highlighted here (e.g. hotspots in Belize, the south of Cuba, northeast of Puerto Rico) mirror recent Caribbean-wide estimations of corals morpho-functional diversity (Melo-Merino et al., 2022), and could indicate priority target for regional conservation initiatives.

*The extended methods and results of this case study are available as supplementary material.*

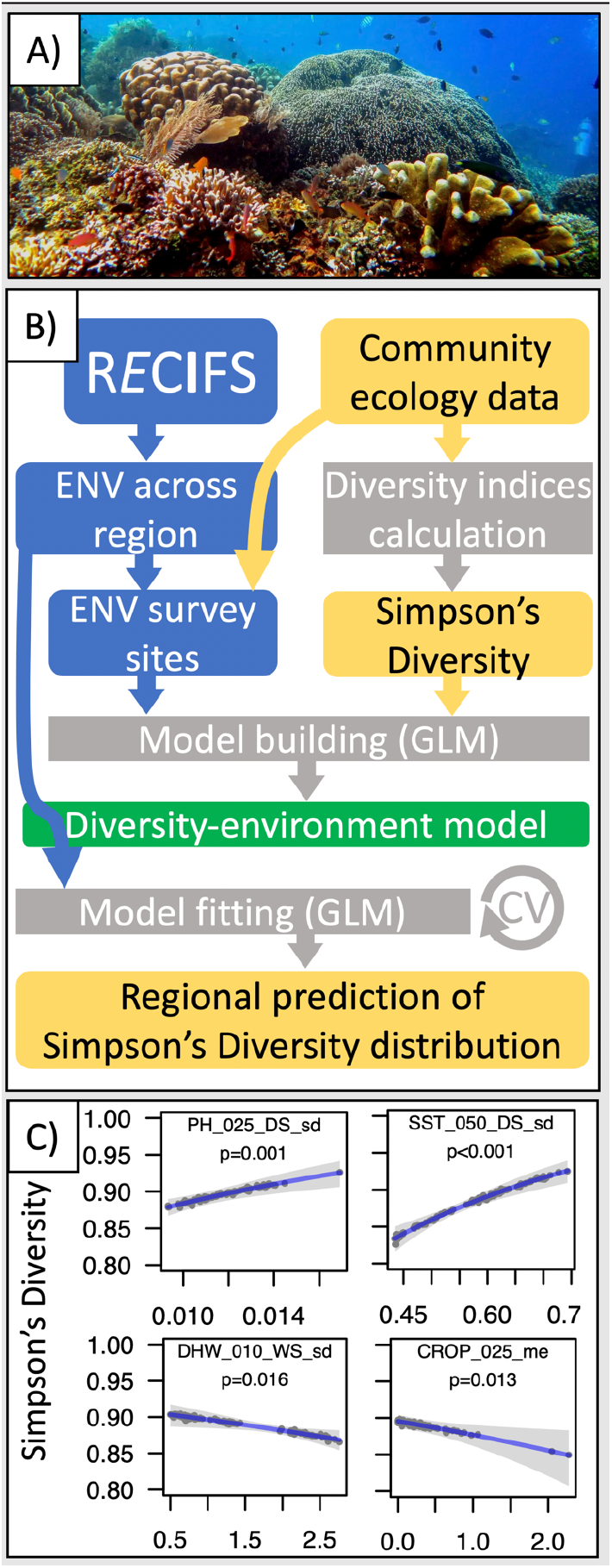
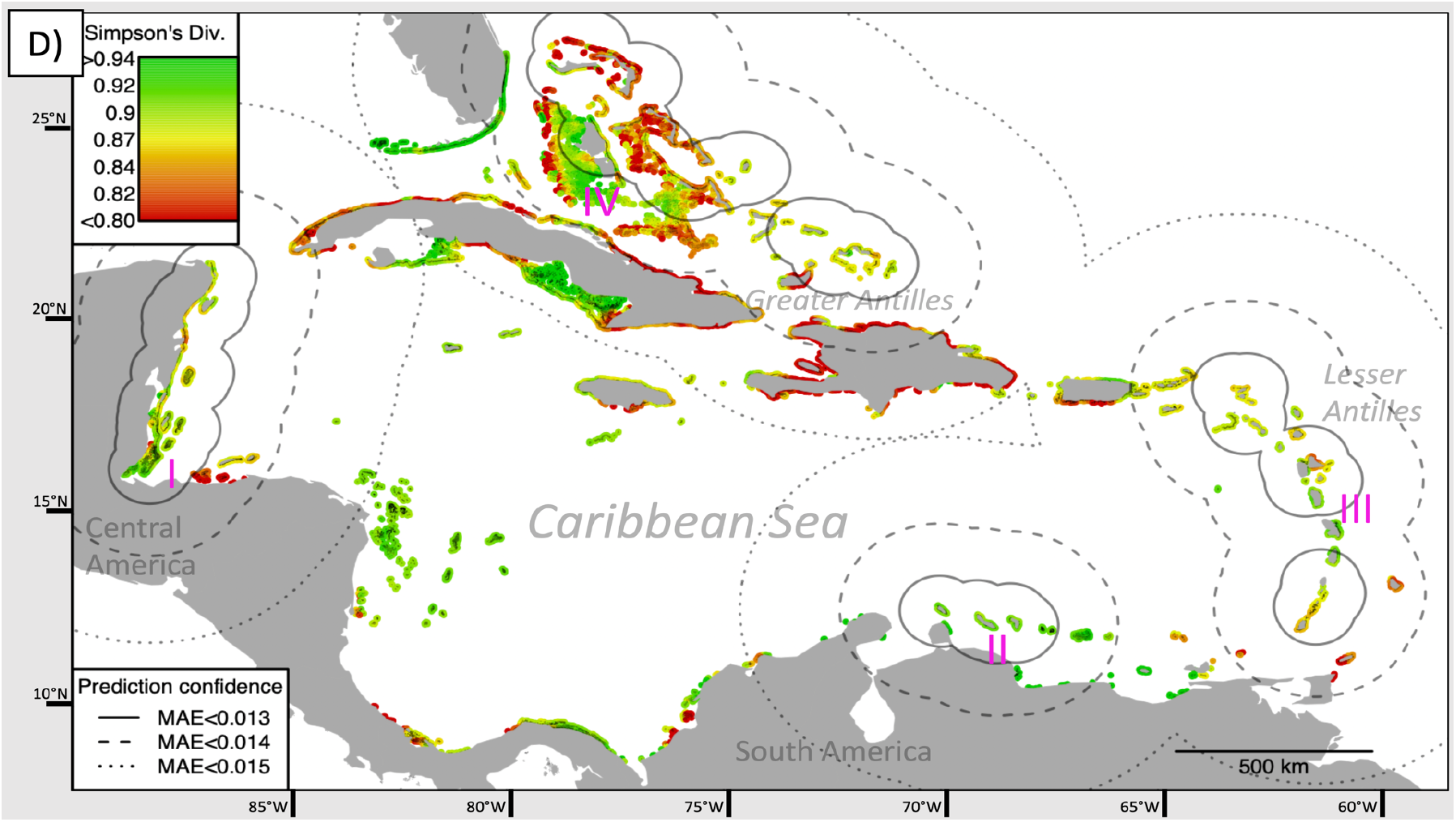

#### Box 2. Stripey snapper adaptation in NW Australia.

For the second case study we use pre-existing population genomics data from 1,016 stripey snapper individuals (*Lutjanus carponotatus*; A) from 51 reefs along the northwestern coast of Australia (DiBattista et al., 2017). Each sample in this dataset was genotyped at 17,007 single nucleotide polymorphisms (SNPs). Using genome scans, we identified 62 outlier SNPs that did not follow the neutral genetic structure of the population (i.e. SNPs potentially involved in local adaptation). We then ran a principal component analysis (PCA) to summarize the allelic frequencies of outlier SNPs into 12 axes of outlier genomic variation (outlier-PCs). Using the analytical pipeline shown in (B), we investigated the association between the spatial distribution of outlier-PCs and 302 variables from RECIFS, characterizing the reef environment for specific spatial and temporal scales.

We ran a redundancy analysis (RDA; C) and found that part of the outlier-PCs variation (R^2^=0.52) could be explained by two environmental variables. The first variable was average Degree Heating Week (DHW_010_OA me), which showed the strongest correlation with the first axis of genomic variation (outlier-PC1). The second environmental variable was the standard deviation of Phosphate Concentration (PO4_002_OA_sd), which was correlated with outlier-PC1 and outlier-PC9. We then performed a functional annotation analysis, and found that outlier-PC1 summarized the variation of SNPs located in genes potentially involved in the detection of temperature stimulus, whereas outlier-PC9 in the immune response.

Finally, we employed the association models from the RDA to predict the spatial distribution of the genetic axes potentially involved in local adaptation (D: left side: outlier-PC1; right side: outlier-PC9). High values of outlier-PC1, indicating a higher frequency of alleles putatively implicated in heat adaptation, were observed in the Shark Bay area (highlighted by the purple “I” on maps). As for outlier-PC9, we assumed that low values corresponded to an adaptation to high variability in Phosphate concentration and observed reef cells with such values in the Shark Bay area and the Kimberly region (“II” on map).

These predictions must interpreted with care, as the mean absolute error (MAE) was non-negligible (∼36% and 14% of the range of outlier-PC1 and outlier-PC9, respectively) and increased with distance from sampling locations. Yet, these results suggest that reefs of the Shark Bay area might host a stripey snapper population with exceptional adaptive capacities to both thermal stress and variability in phosphate concentration, compared with the reefs in the northeastern regions. In the original work that produced the genomic dataset used in this case study, DiBattista and colleagues highlighted a connectivity break between the reefs in Shark Bay and those located in northern areas (DiBattista et al., 2017). For these reasons, stripey snapper in the Shark Bay area might be of particular interest for conservation efforts.

*The extended methods and results of this case study are available as supplementary material.*

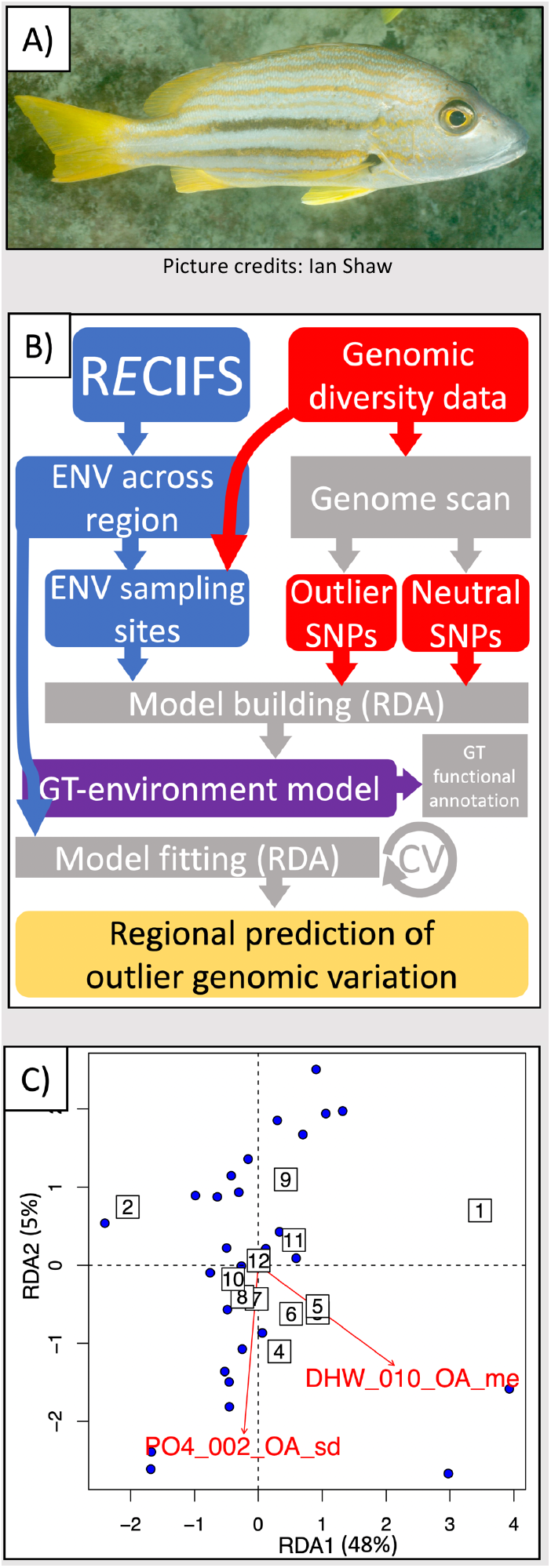
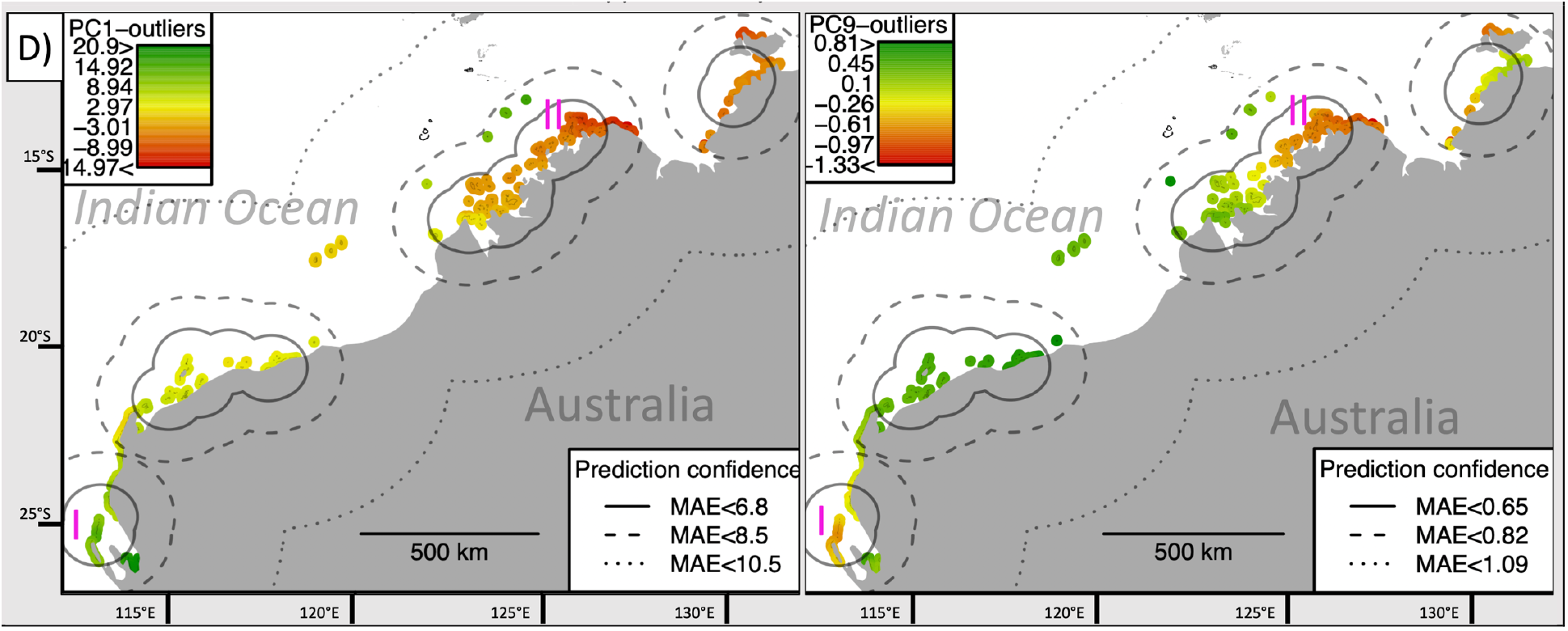

## Acknowledgments

We thank the environmental data providers for sharing the datasets included in RECIFS. We thank the Catlin Seaview Project and Joseph DiBattista for sharing the data used for the case studies. We thank Annie Guillaume for the comments and suggestions provided during the redaction of this paper. The authors declare no conflict of interest.

## Notes

https://recifs.epfl.ch

### Competing Interest Statement

The authors have declared no competing interest.

