## Supplementary Material for "RECIFS: a centralized geo-environmental database for coral reef research and conservation"

### Extended Methods

We present two case studies to illustrate the use of the RECIFS tool (Supplementary Figure 1). The first investigates the environmental drivers of coral taxonomic diversity in the Caribbean, and the second explores the environmental factors shaping the adaptive genomic diversity of a reef fish (stripey snapper).

**Supplementary Figure 1. Case studies.** The first case study investigates the environmental drivers of coral taxonomic diversity in the Caribbean (A-C). The data used in this analysis comes from field surveys performed across 183 reefs (red points in B) by the Catlin Seaview Survey project (González-Rivero et al., 2014, 2016). The second case study explores the environmental factors shaping the adaptive genomic diversity of a reef fish (stripey snapper) in Northwest Australia (D-F). The genomic data used in this analysis originate from a previous population genomics study across 51 reefs of the region (red points in E; DiBattista et al., 2017). The diagrams in C) and F) outline the analytical pipeline followed in the coral diversity and fish adaptation study, respectively. (credit for picture D: Ian Shaw)

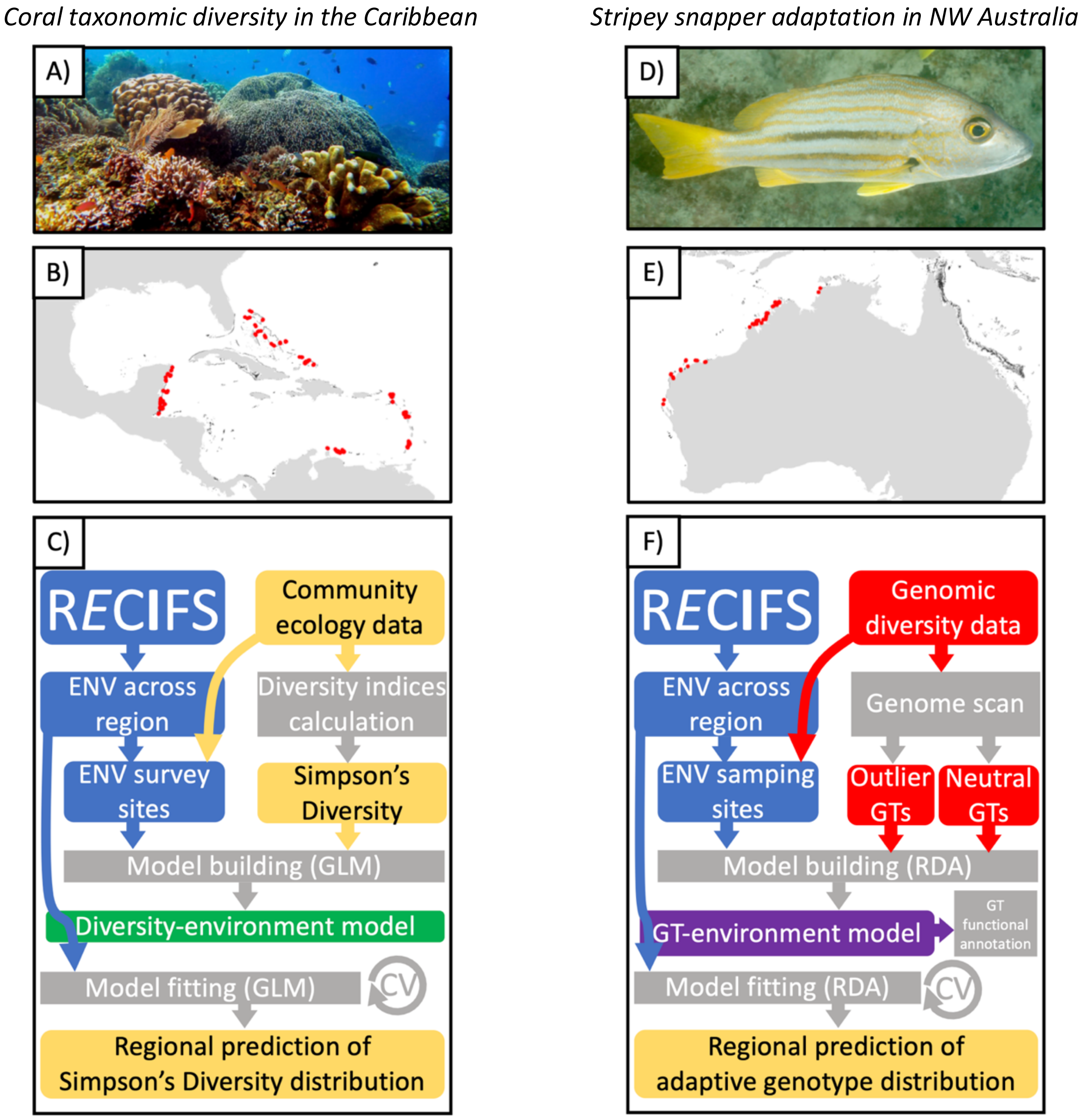

*Case study 1: Coral taxonomic diversity in the Caribbean*

The first case study was based on ecological survey data collected across the reefs of the Caribbean by the Catlin Seaview Survey (CSS) project (Supplementary Figure 1A-C; González-Rivero et al., 2014, 2016). This dataset included 183 field surveys performed in 2013 across the reefs of 12 territories of the region (Anguilla, Aruba, Bahamas, Belize, Bonaire, Curaçao, Guadeloupe, Mexico, St. Martin, St. Vincent and the Grenadines, Turks and Caicos, United States of America; Supplementary Figure 1B). Each survey consisted in a photo-transect along ~1.5-2 km of reef at a constant depth of 10 meters. The CSS project processed survey pictures using machine learning trained to recognize 17 hard coral taxa manually labelled by coral taxonomists (Beijbom et al., 2012). The taxon-labels referred to species (e.g. *Acropora cervicornis, Montastraea cavernosa, Porites astreoides*) or to groups of species with similar morphology (e.g. *Porites* branching, *Orcbicella* complex; Supplementary Table 1).

We retrieved the coordinates of each survey location and grouped together surveys located less then 5 km apart (based on Euclidean distance of the geodetic coordinates, calculated used the SoDA R package, v 1.0; Chambers, 2020). This resulted in 124 groups of surveys, hereafter referred to as "survey units". For each survey unit, we calculated the abundance of corals from each taxon (i.e. the percentage of pictures in which the corresponding taxon-label was observed). The result was a community matrix with survey units as rows and taxa abundance as columns. Using the R vegan package (v.2.5; Oksanen et al., 2019), we applied a Hellinger normalization (Legendre & Gallagher, 2001) and then calculated Simpson's diversity index (SDI) for every survey unit (Simpson, 1949).

We used RECIFS to then characterize the environment of the reef system of the Caribbean. Such characterization was based the "Advanced environment query" targeting the "Extended tables" of RECIFS, and differed between variables with different temporal resolutions. For variables with monthly temporal resolution, we focused on records before 2013 and computed mean and standard deviation for (1) all months, (2) months in wet season (May to November) and (3) months in the dry season (December to April). For variables with annual temporal resolutions, we computed mean and standard deviation for all the years available, whereas for variables with no temporal resolution (bathymetry and surface land) no further calculations were performed. All of these calculations were replicated for the four different buffer sizes (2.5, 10, 25 and 50 km), resulting in a total of 304 environmental variables characterizing the reef environment. Missing values were generally infrequent (median frequency: 0% [0%-10%] of the reef cells) and were imputed using a random forest approach, as featured in the MissForest R package (v. 1.4; Stekhoven & Bühlmann, 2012). The imputation error (calculated using the built-in "Out-of-bag" method) was estimated to have a median magnitude of 0.05% [0%-0.1%] of the variable's standard deviation (Supplementary Table 2a). Only one variable displayed higher levels of imputation error (bathymetry at 2.5 km buffer, with an error estimated to more than 100% of the variable's standard deviation) and was excluded from downstream analyses. For each survey unit, environmental values were assigned from those of the closest reef cell from RECIFS.

We then ran an association study between SDI and the RECIFS environmental data. This was done by constructing a generalized linear model (GLM) where SDI was the logit-transformed response variable. As for the explanatory variables, we first created groups of highly-collinear environmental variables ($R>0.8$) and randomly picked one variable per group for an initial stage of model selection. Using the MASS R package (v. 7.3; Venables & Ripley, 2002), we ran a stepwise model selection (based on the Akaike Information Criterion, AIC; Bozdogan, 1987) for the resulting 35 non-collinear environmental variables, with the goal of identifying the most parsimonious model. This model retained eleven environmental variables. We finalized the model construction by checking whether any of these eleven variables could be replaced by a highly-collinear variable maximizing the quality-of-fit, according to AIC. The explanatory variables retained in this final model were evaluated by observing their effects, the associated T-statistics and the corresponding p-values.

We used the final model to predict SDI from the RECIFS environmental data, and to represent such predictions on a map for every reef cell of the Caribbean. To assess the predictive power of this approach, we ran a spatially-explicit K-fold cross validation (Pohjankukka et al., 2017), with the following procedure:

(1) create geographical clusters of survey units that are located within a distance *D* from each other.

(2a) train the final model leaving one geographical cluster out.

(2b) use the model trained in (2a) to predict SDI for the geographical cluster not used in model training.

(3) repeat step (2a-b) for each geographical cluster.

(4) calculate the mean absolute error (MAE) of the difference between real and predicted SDI values.

These four steps were replicated for different sizes of *D* (25, 50, 100, 250 and 500 km), with each replicate resulting in a specific MAE. These estimations of error were then added to the map of the SDI predictions as isolines surrounding the survey units at distance *D*.

*Case study 2: Stripey snapper adaptation in NW Australia*

For the second case study, we used pre-existing population genomics data on the stripey snapper (*Lutjanus carponotatus*; Supplementary Figure 1D) from the reefs along the northwestern coast of Australia (DiBattista et al., 2017). This dataset was produced for a connectivity study where 1,016 fish were sampled from 51 reefs (Supplementary Figure 1E) of the area between 2014 and 2015. Samples in this dataset were genotyped at 17,007 single nucleotide polymorphisms (SNPs) using a restriction-sites-associated DNA sequencing (RADseq) method - called Diversity Array Technology sequencing (DArT-seq) – targeting the functional regions of the genome (e.g. genes, regulatory sequences; Andrews et al., 2016; Kilian et al., 2012; Sansaloni et al., 2011).

We used the R package poppr (v. 2.9; Kamvar et al., 2014, 2015) to filter out (1) samples with excessively high number of missing SNPs, (2) SNPs missing in excessively high number of individuals, and (3) SNPs with excessively low frequencies of minor alleles (the cut-off threshold was set to 5% for every filter). After these filtering steps, 5,971 SNPs across 671 samples were retained for downstream analyses. We then used the PCadapt R package (v. 4.3; Luu et al., 2017) and identified 62 outlier SNPs, *i.e.* SNPs whose distribution deviates from the neutral genetic distribution across the population, therefore possibly implicated in adaptive processes.

Using the R SoDA package, we grouped together samples collected less then 5 km apart into 37 sampling units, and for each of these units we calculated the allelic frequency of each SNP. To reduce bias in the estimation of genetic frequencies per sampling unit, the analysis focused on sampling units (28 out of 37) with at least 10 samples. The results were two matrices describing allelic frequencies across 28 sampling units: one for the 62 outlier loci and one for the 5,908 neutral (non-outlier) loci. Each matrix was normalized using the Hellinger method (Legendre & Gallagher, 2001) and then summarized running a Principal Component Analysis

(PCA; Novembre et al., 2008) to retain only the Principal Component axes (PCs) cumulatively explaining up to 90% of the overall allelic variance (19 neutral-PCs and 12 outlier-PCs).
The environmental data across the reefs of the region were extracted from RECIFS and then processed using the same methods from Case Study 1. In short, we computed means and standard deviations both overall and by season (wet season: December to March, dry season: April to November) for environmental variables measured at monthly temporal resolution. For variables at yearly temporal resolution, we only computed overall mean and standard deviation. Next, we used the MissForest algorithm to impute missing environmental data (median frequency of missing data: 0% [0%-5%]; median magnitude of imputation error: 0.06% [0%-0.2%] of the variable's standard deviation; Supplementary Table 2b). For each sampling unit, environmental values were attributed from the closest reef cell from RECIFS.
The redundancy analysis (RDA) was then run to investigate the association between genetic variation of the stripey snapper and environmental variation of the reefs of the region. Using the vegan R package (v.2.5; Oksanen et al., 2019), the RDA was performed in two steps: we first characterized the spatial structure of neutral genetic variation; and then investigated the association between environmental variables and outlier genetic variation, while accounting for such neutral spatial structure.
We started by computing (pcnm function in vegan) the distance-based Moran's Eigenvector Maps (dbMEMs), which are a matrix of orthogonal vectors synthesizing the spatial distances between sampling units (Borcard & Legendre, 2002). We ran a stepwise RDA (ordistep function with 999 permutations) to uncover which dbMEMs explained the variation of the neutral-PCs, *i.e.* the dbMEMs retained in the most parsimonious RDA model according to the coefficient of determination ($R^2$). Only the first dbMEM was retained in this model ($R^2$=0.05).
Next, we performed a stepwise RDA (999 permutations), this time to investigate the association between outlier-PCs environmental variables from RECIFS. This RDA was run as a partial-RDA, where the first dbMEM was employed as a conditional variable to control for neutral spatial structure of genetic variation. Similar to case study 1, we first performed model selection using 32 (randomly selected) non-collinear (R<0.8) environmental variables to identify the most parsimonious model. Next, we built the final model by replacing the retained environmental variables with highly-collinear predictors maximizing the $R^2$ coefficient. The effect of the predictors retained in the final model was assessed using an analysis of variance (ANOVA; 999 permutations).
Finally, we investigated which outlier-PCs appeared to be more strongly associated with the environmental variables retained in the final model by: (1) manually inspecting the RDA tri-plot and (2) checking the correlation between outlier-PCs and environmental variables. Such outlier-PCs might represent axes of adaptive genetic variation, and we used the final model to predict their spatial distribution for every reef cell of NW Australia. Using the same method described in case study 1, we then ran a spatially-explicit K-fold cross validation of such predictions.

### Extended Results & Discussion

*Case study 1: Coral taxonomic diversity in the Caribbean*

We conducted two case studies to show how the data from RECIFS can be used to characterize the environmental factors shaping the dynamics of the coral reef ecosystem.
The first case study investigated the environmental drivers of hard coral diversity across the reefs of the Caribbean. Using RECIFS, we produced 302 variables - with varying spatial and temporal resolutions - describing the environmental variation across the reefs of the area. We found 11 of these variables significantly ($p<0.05$) explained more than half of variation ($R^2=0.56$; Supplementary Figure 2A) of the Simpson's Diversity Index (SDI), which summarized the abundance of 17 coral taxa (González-Rivero et al., 2014, 2016). These 11 variables referred to environmental conditions well known to mediate changes in coral abundance/mortality, such as thermal stress (standard deviation of Degree Heating Week, calculated during the wet season using a buffer of 10 km), ocean acidification (standard deviation of pH, dry season, 25 km buffer), water turbidity (average Suspended matter concentration, wet season, 10 km buffer) or run-off from agricultural activities along the coastline (land use of cropland, 25 km buffer; Cornwall et al., 2021; D'Angelo & Wiedenmann, 2014; De'ath & Fabricius, 2010; R. Jones et al., 2020; McClanahan et al., 2020; Sully et al., 2019). However, interpreting how a given variable might be responsible for an increase or a decrease in SDI is not straightforward, mainly because different variables relating to a same condition (e.g. standard deviation vs. mean of Degree Heating Week) are often highly collinear, which makes hard to disentangle their effects on SDI (Supplementary Figure 3).
Based on the associations described here above, the RECIFS data was employed to predict SDI for every reef cell of the Caribbean (Supplementary Figure 2B). This spatial prediction highlighted regions expected to host reefs with high coral diversity, for instance in Belize (South of zone A); Bonaire, Aruba and Curaçao (zone D); Dominica and Martinique (center of zone E); and the Andros Island in Bahamas (Southwest of zone B). The mean absolute error of this prediction was 0.013±0.012 (i.e. ~11% of the range of SDI measured in real surveys), and did not appear to increase with distance from survey locations (Supplementary Figure 4). Yet, care must be taken when considering this prediction since the survey data was systematically collected at 10 meters of depth, and these diversity estimations might therefore not be relevant at other depth levels. Some of the biodiversity patterns highlighted here (*e.g.* hotspots in Belize, the south of Cuba, northeast of Puerto Rico) mirror recent Caribbean-wide estimations of corals morpho-functional diversity (Melo-Merino et al., 2022), and could indicate priority target for regional conservation initiatives.

**Supplementary Figure 2. Case study 1: Coral taxonomic diversity in the Caribbean.** An association model was built to describe the link between Simpson Diversity Index (SDI) from coral field surveys and 302 environmental variables produced using from RECIFS. Eleven variables were retained in such model, and their associations with SDI are shown in A (blue regression line, with the grey band showing the 95% interval of confidence). The identifier of the environmental variable involved is shown on top of every plot, and such identifier is defined by (1) the name of the environmental descriptor (DHW: degree heating week, FE: iron concentration, O2: oxygen concentration, PH: water pH, PO4: phosphate concertation, SPM: suspended matter concentration, SST: sea surface temperature, CROP: density of cropland areas, LAND: density of surface land); followed by (2) the buffer size from the survey site, used to calculate the spatial resolution of the variable (002: 2.5 km, 010: 10 km, 025: 25 km and 050: 50 km); followed by (3) the temporal window during which the variable was calculated (DS: dry season, WS: wet season, OA: overall); followed by (4) the statistic used to summarize the environmental variable over time (me: mean, sd: standard deviation). For every plot, the p-value of the T-statistic describing the association is shown below the identifier.
This association model was then used to predict SDI for every reef cell (points) of the Caribbean (B). The isolines on the map display the spatial boundaries of three different levels of confidence in the prediction, each defined by a corresponding Mean Absolute Error (MAE). The map details on the right show the real SDI values of the surveys (circled points) used to train the model.

A)

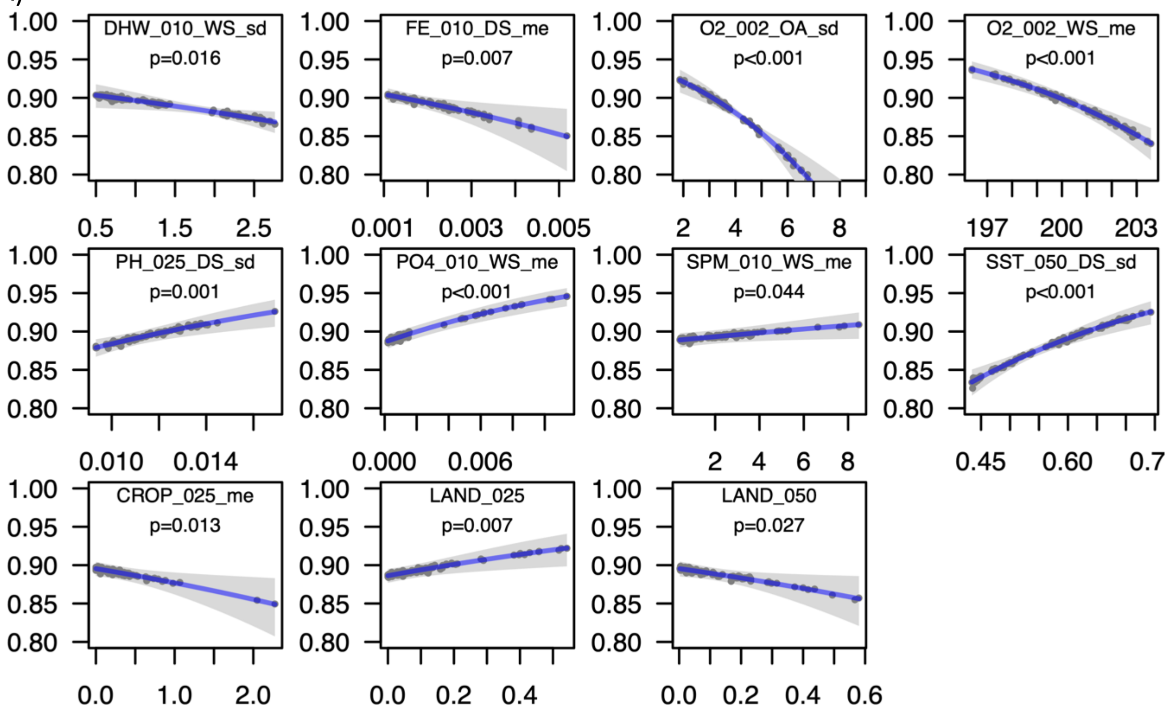


B)

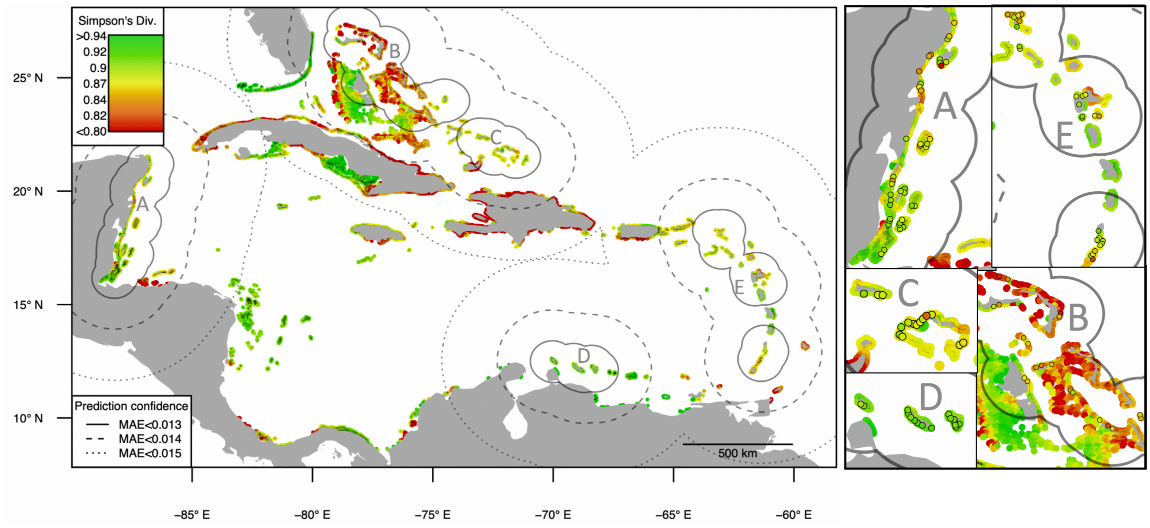

**Supplementary Figure 3. Correlation between environmental variables in case study 1.** The plot displays the levels of average correlation between 35 groups of highly correlated environmental variables (clsXX). The diagonal indicates the average correlation within each group. Below the plot, the variables included in every group of correlated variables are listed. The identifier of the environmental variable involved is shown on top of every plot, and such identifier is defined by (1) the name of the environmental descriptor (CHL: chlorophyll concentration, DHW: degree heating week, FE: iron concentration, NO3: nitrate concentration, O2: oxygen concentration, PH: water pH, PO4: phosphate concertation, SPM: suspended matter concentration, SSS: sea surface salinity, SST: sea surface temperature, CROP: density of cropland areas, LAND: density of surface land, URBA: density of built-up areas, VBD: average boat detection, PDEN: human population density); followed by (2) the buffer size from the survey site, used to calculate the spatial resolution of the variable (002: 2.5 km, 010: 10 km, 025: 25 km and 050: 50 km); followed by (3) the temporal window during which the variable was calculated (DS: dry season, WS: wet season, OA: overall); followed by (4) the statistic used to summarize the environmental variable over time (me: mean, sd: standard deviation).

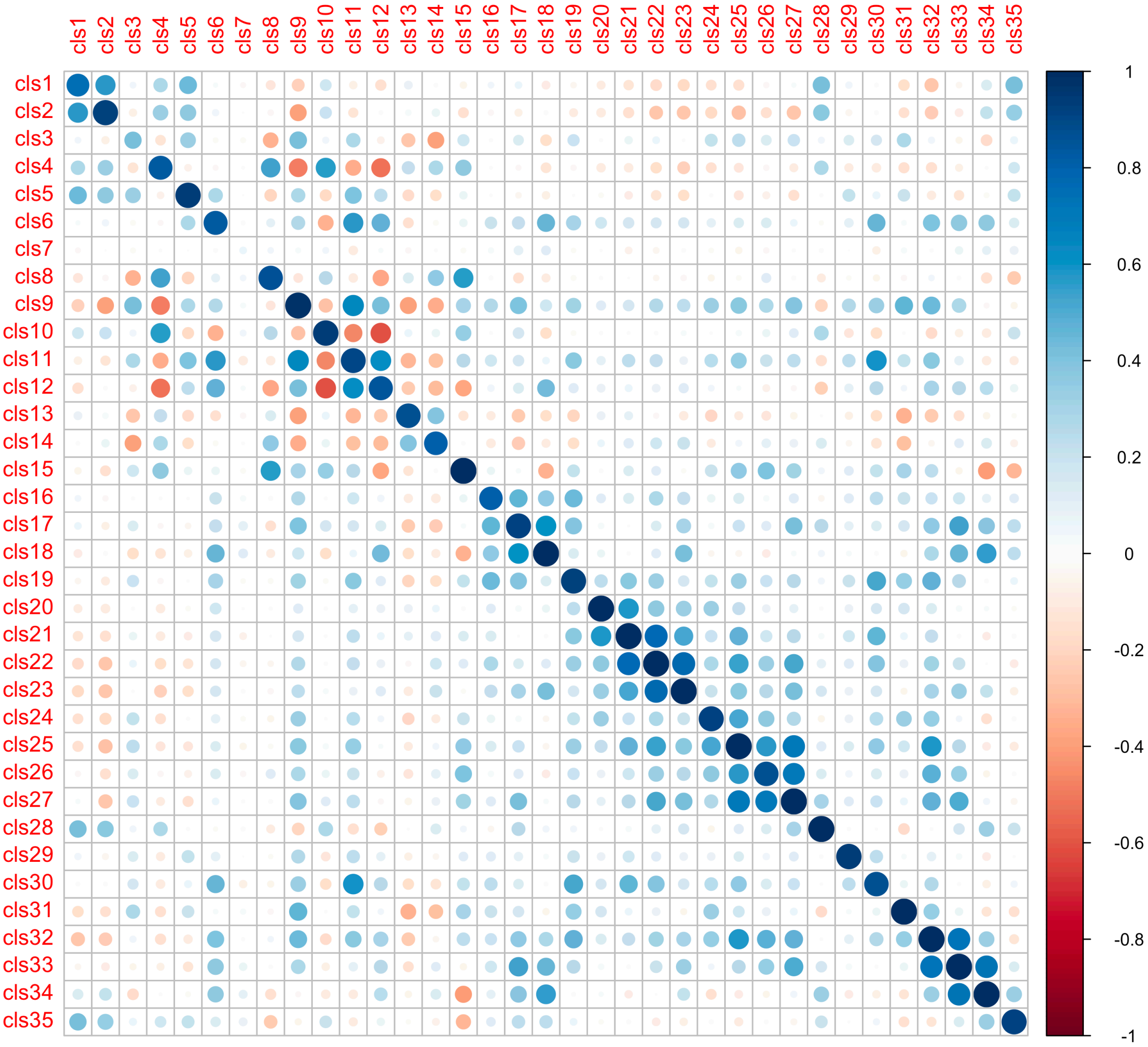


| | |
|---|---|
| **cls1** | CHL_002_OA_me, CHL_002_OA_sd, CHL_002_WS_me, CHL_002_WS_sd, CHL_002_DS_me, CHL_002_DS_sd, CHL_010_OA_me, CHL_010_OA_sd, CHL_010_WS_me, CHL_010_WS_sd, CHL_010_DS_me, CHL_010_DS_sd, CHL_025_OA_me, CHL_025_OA_sd, CHL_025_WS_me, CHL_025_WS_sd, CHL_025_DS_me, CHL_025_DS_sd |
| **cls2** | CHL_050_OA_me, CHL_050_OA_sd, CHL_050_WS_me, CHL_050_WS_sd, CHL_050_DS_me, CHL_050_DS_sd |
| **cls3** | DHW_002_OA_me, DHW_002_OA_sd, DHW_002_WS_me, DHW_002_WS_sd, DHW_002_DS_me, DHW_002_DS_sd, DHW_010_OA_me, DHW_010_OA_sd, DHW_010_WS_me, DHW_010_WS_sd, DHW_010_DS_me, DHW_010_DS_sd, DHW_025_OA_me, DHW_025_OA_sd, DHW_025_WS_me, DHW_025_WS_sd, DHW_025_DS_me, DHW_025_DS_sd, DHW_050_OA_me, DHW_050_OA_sd, DHW_050_WS_me, DHW_050_WS_sd, DHW_050_DS_me, DHW_050_DS_sd, PH_002_OA_me, PH_002_WS_me, PH_002_DS_me, PH_010_OA_me, PH_010_WS_me, PH_010_DS_me, PH_025_OA_me, PH_025_WS_me, PH_025_DS_me, PH_050_OA_me, PH_050_WS_me, PH_050_DS_me, SSS_002_OA_me, SSS_002_OA_sd, SSS_002_WS_me, SSS_002_WS_sd, SSS_002_DS_me, SSS_002_DS_sd, SSS_010_OA_me, SSS_010_OA_sd, SSS_010_WS_me, SSS_010_WS_sd, SSS_010_DS_me, SSS_010_DS_sd, SSS_025_OA_me, SSS_025_OA_sd, SSS_025_WS_me, SSS_025_WS_sd, SSS_025_DS_me, SSS_025_DS_sd, SSS_050_OA_me, SSS_050_OA_sd, SSS_050_WS_me, SSS_050_WS_sd, SSS_050_DS_me, SSS_050_DS_sd |
| **cls4** | FE_002_OA_me, FE_002_OA_sd, FE_002_WS_me, FE_002_WS_sd, FE_002_DS_me, FE_002_DS_sd, FE_010_OA_me, FE_010_OA_sd, FE_010_WS_me, FE_010_WS_sd, FE_010_DS_me, FE_010_DS_sd, FE_025_OA_me, FE_025_OA_sd, FE_025_WS_me, FE_025_WS_sd, FE_025_DS_me, FE_025_DS_sd, FE_050_OA_me, FE_050_OA_sd, FE_050_WS_me, FE_050_WS_sd, FE_050_DS_me, FE_050_DS_sd |
| **cls5** | NO3_002_OA_me, NO3_002_OA_sd, NO3_002_WS_me, NO3_002_WS_sd, NO3_010_OA_me, NO3_010_OA_sd, NO3_010_WS_me, NO3_010_WS_sd, NO3_025_OA_me, NO3_025_OA_sd, NO3_025_WS_me, NO3_025_WS_sd, NO3_050_OA_me, NO3_050_OA_sd, NO3_050_WS_me, NO3_050_WS_sd |
| **cls6** | NO3_002_DS_me, NO3_002_DS_sd, NO3_010_DS_me, NO3_010_DS_sd, NO3_025_DS_me, NO3_025_DS_sd, NO3_050_DS_me, NO3_050_DS_sd |
| **cls7** | O2_002_OA_me, O2_010_OA_me, O2_025_OA_me, O2_050_OA_me, SST_002_WS_me, SST_010_WS_me, SST_025_WS_me, SST_050_WS_me |
| **cls8** | O2_002_OA_sd, O2_002_WS_sd, O2_002_DS_me, O2_002_DS_sd, O2_010_OA_sd, O2_010_WS_sd, O2_010_DS_me, O2_010_DS_sd, O2_025_OA_sd, O2_025_WS_sd, O2_025_DS_me, O2_025_DS_sd, O2_050_OA_sd, O2_050_WS_sd, O2_050_DS_me, O2_050_DS_sd, PH_002_OA_sd, PH_002_WS_sd, PH_010_OA_sd, PH_010_WS_sd, PH_025_OA_sd, PH_025_WS_sd, PH_050_OA_sd, PH_050_WS_sd, SST_002_OA_me, SST_002_OA_sd, SST_002_WS_sd, SST_002_DS_me, SST_010_OA_me, SST_010_OA_sd, SST_010_WS_sd, SST_010_DS_me, SST_025_OA_me, SST_025_OA_sd, SST_025_WS_sd, SST_025_DS_me, SST_050_OA_me, SST_050_OA_sd, SST_050_WS_sd, SST_050_DS_me |

| | |
|---|---|
| **cls9** | O2_002_WS_me, O2_010_WS_me, O2_025_WS_me, O2_050_WS_me |
| **cls10** | PH_002_DS_sd, PH_010_DS_sd, PH_025_DS_sd, PH_050_DS_sd |
| **cls11** | PO4_002_OA_me, PO4_002_OA_sd, PO4_002_WS_me, PO4_002_WS_sd, PO4_002_DS_me, PO4_002_DS_sd, PO4_010_OA_me, PO4_010_OA_sd, PO4_010_WS_me, PO4_010_WS_sd, PO4_010_DS_me, PO4_010_DS_sd, PO4_025_OA_me, PO4_025_OA_sd, PO4_025_WS_me, PO4_025_WS_sd, PO4_025_DS_me, PO4_025_DS_sd, PO4_050_OA_me, PO4_050_OA_sd, PO4_050_WS_me, PO4_050_WS_sd, PO4_050_DS_me, PO4_050_DS_sd |
| **cls12** | SCV_002_OA_me, SCV_002_OA_sd, SCV_002_WS_me, SCV_002_WS_sd, SCV_002_DS_me, SCV_002_DS_sd, SCV_010_OA_me, SCV_010_OA_sd, SCV_010_WS_me, SCV_010_WS_sd, SCV_010_DS_me, SCV_010_DS_sd, SCV_025_OA_me, SCV_025_OA_sd, SCV_025_WS_me, SCV_025_WS_sd, SCV_025_DS_me, SCV_025_DS_sd, SCV_050_OA_me, SCV_050_OA_sd, SCV_050_WS_me, SCV_050_WS_sd, SCV_050_DS_me, SCV_050_DS_sd |
| **cls13** | SPM_002_OA_me, SPM_002_OA_sd, SPM_002_WS_me, SPM_002_WS_sd, SPM_002_DS_me, SPM_002_DS_sd |
| **cls14** | SPM_010_OA_me, SPM_010_OA_sd, SPM_010_WS_me, SPM_010_WS_sd, SPM_010_DS_me, SPM_010_DS_sd, SPM_025_OA_me, SPM_025_OA_sd, SPM_025_WS_me, SPM_025_WS_sd, SPM_025_DS_me, SPM_025_DS_sd, SPM_050_OA_me, SPM_050_OA_sd, SPM_050_WS_me, SPM_050_WS_sd, SPM_050_DS_me, SPM_050_DS_sd |
| **cls15** | SST_002_DS_sd, SST_010_DS_sd, SST_025_DS_sd, SST_050_DS_sd |
| **cls16** | PDEN_002_me, PDEN_002_sd, PDEN_010_me, PDEN_010_sd, PDEN_025_me, PDEN_025_sd, URBA_025_me |
| **cls17** | PDEN_050_me, URBA_050_me |
| **cls18** | PDEN_050_sd |
| **cls19** | URBA_002_me, URBA_010_me |
| **cls20** | URBA_002_sd |
| **cls21** | URBA_010_sd |
| **cls22** | URBA_025_sd |
| **cls23** | URBA_050_sd |
| **cls24** | CROP_002_me, CROP_002_sd |
| **cls25** | CROP_010_me |
| **cls26** | CROP_010_sd, CROP_025_sd, CROP_050_sd |
| **cls27** | CROP_025_me |
| **cls28** | CROP_050_me |
| **cls29** | VBD_002_me, VBD_002_sd |
| **cls30** | VBD_010_me, VBD_010_sd, VBD_025_me, VBD_025_sd, VBD_050_me, VBD_050_sd |
| **cls31** | LAND_002 |
| **cls32** | LAND_010 |
| **cls33** | LAND_025 |
| **cls34** | LAND_050 |
| **cls35** | BATHY_010, BATHY_025, BATHY_050 |

**Supplementary Figure 4. Absolute error from spatially-explicit cross validations of case study 1.** The boxplot displays the distribution of the absolute error of the Simpsons Diversity Index predicted by the cross-validation procedure. Each box corresponds to the iterations performed under a given value of the threshold distance D, which was used to group together survey sites that are spatially close in the spatially-explicit cross validation procedure.

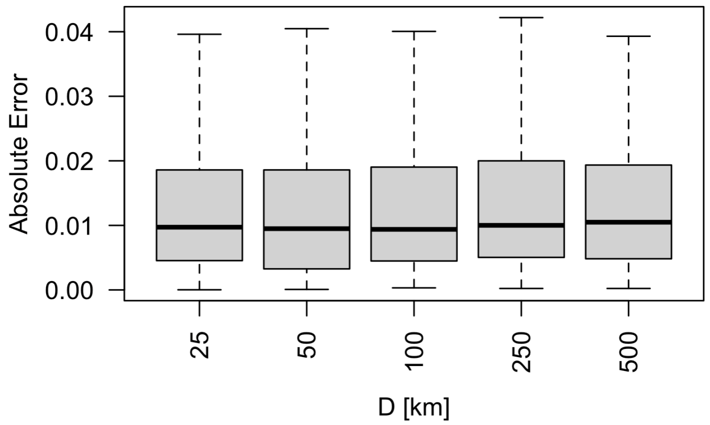

*Case study 2: Stripey snapper adaptation in NW Australia*

The second case study investigated how environmental variability shapes genetic diversity - potentially driving local adaptation - of a reef fish population (stripey snapper) from the northwestern coast of Australia (DiBattista et al., 2017). We identified 62 outlier single nucleotide polymorphisms (SNPs) not following the neutral genetic structure of the population, and summarized their allelic frequencies in 12 axes of genomic variation (outlier-PCs). The redundancy analysis (RDA) revealed that two environmental variables (out of 302) from RECIFS that explained part of the variation across these genomics axes (R2=0.52, Supplementary Figure 5A). The first variable was average Degree Heating Week (DHW, calculated using a 10 km buffer across all the months of the year; p=0.001 under ANOVA), which showed the strongest correlation (R=0.65) with the first axis of genomic variation (outlier-PC1; Supplementary Figure 5B). DHW is a coral bleaching-related indicator of thermal stress (Liu et al., 2003), where here the variable could also act as a generic proxy of thermal stress for stripey snapper. To this hypothesis, we performed a functional annotation analysis suggesting that outlier-PC1 summarized the variation of SNPs located in genes involved in the biological process "Detection of temperature stimulus" (Supplementary Box 1). The second environmental variable was the standard deviation of Phosphate Concentration (2 km buffer, across all the months of the year; p=0.006 under ANOVA), which was correlated with outlier-PC1 (R=0.65) and outlier-PC9 (R=0.40; Supplementary Figure 5B). The latter summarized the variation of SNPs located in genes potentially involved in the fish immune response (Suppelementary Box 1). The effects of dietary phosphorus intake on the immune system has already been highlighted in freshwater fish (Wang et al., 2017; Yang et al., 2021), and should be further investigated in stripey snapper.
Like in the first case study, we employed the association models from the RDA to predict the spatial distribution of the genetic axes potentially involved in local adaptation (outlier-PC1, Supplementary Figure 5C; outlier-PC9, Supplementary Figure 5D). High values of outlier-PC1, indicating a higher frequency of alleles putatively implicated in heat adaptation, were observed in the Shark Bay area (zone A in Supplementary Figure 5C). As for outlier-PC9, we assumed that low values corresponded to an adaptation to high variability in Phosphate concentration, and observed reef cells with such values in the Shark bay area (zone A in Supplementary Figure 5D) and the Kimberly region (North of zone C). The result of such predictions must be taken with care, since the mean absolute error was non-negligible (~36% and to 14% of the range of outlier-PC1 and outlier-PC9, respectively) and increased along with distance from sampling locations (Supplementary Figure 7). If confirmed by further assays (e.g. eco-physiological studies and gene-expression experiments), these predictions suggest that reefs of the Shark Bay area might host a stripey snapper population with exceptional adaptive capacities to thermal stress and variability in phosphate concentration, compared with the reefs in the northeastern regions. In the original work that produced the genomic dataset used in this case study, DiBattista and colleagues highlighted a connectivity break between the Shark Bay area and those from the reefs located North (DiBattista et al., 2017). For all these reasons, stripey snapper in the Shark Bay area might be of particular interest for conservation efforts.

**Supplementary Figure 5. Case study 2: Stripey snapper adaptation in NW Australia.** A population genomic study on stripey snapper from the reefs of the northwestern Australia revealed 62 single nucleotide polymorphisms (SNPs) not following the neutral genetic structure of the population (i.e. outliers). We ran a redundancy analysis (RDA) to investigate how the axes of variation of these SNPs (outlier-PCs) were associated with environmental variables from RECIFS (A). We found two environmental variables significantly explaining ($p<0.01$ under analysis of variance) allelic variation across two redundancy axes (explaining 48% and 5% of the overall variation): (1) DHW_010_OA_me (overall mean of degree heating week, measured in a 10 km buffer around sampling sites), (2) PO4_002_OA_sd (overall standard deviation of phosphate concentration measured in a 2.5 km buffer). The colors on the RDA tri-plot indicate groups of sampling site close to each other, as shown on the map on the right. (B) shows the correlation between these two environmental variables and the 12 outlier-PCs. The RDA model was used to predict the spatial distribution of outlier-PCs for every reef of the study area. We display the map of such prediction for two outlier-PCs strongly correlated with the environmental variables retained in the RDA: outlier-PC1 (C) and outlier-PC9 (D). The isolines on the map display the spatial boundaries of three different levels of confidence in the prediction, each defined by a Mean Absolute Error (MAE). The map details on the right show the real outlier-PC values of the surveys (circled points) used to train the model.

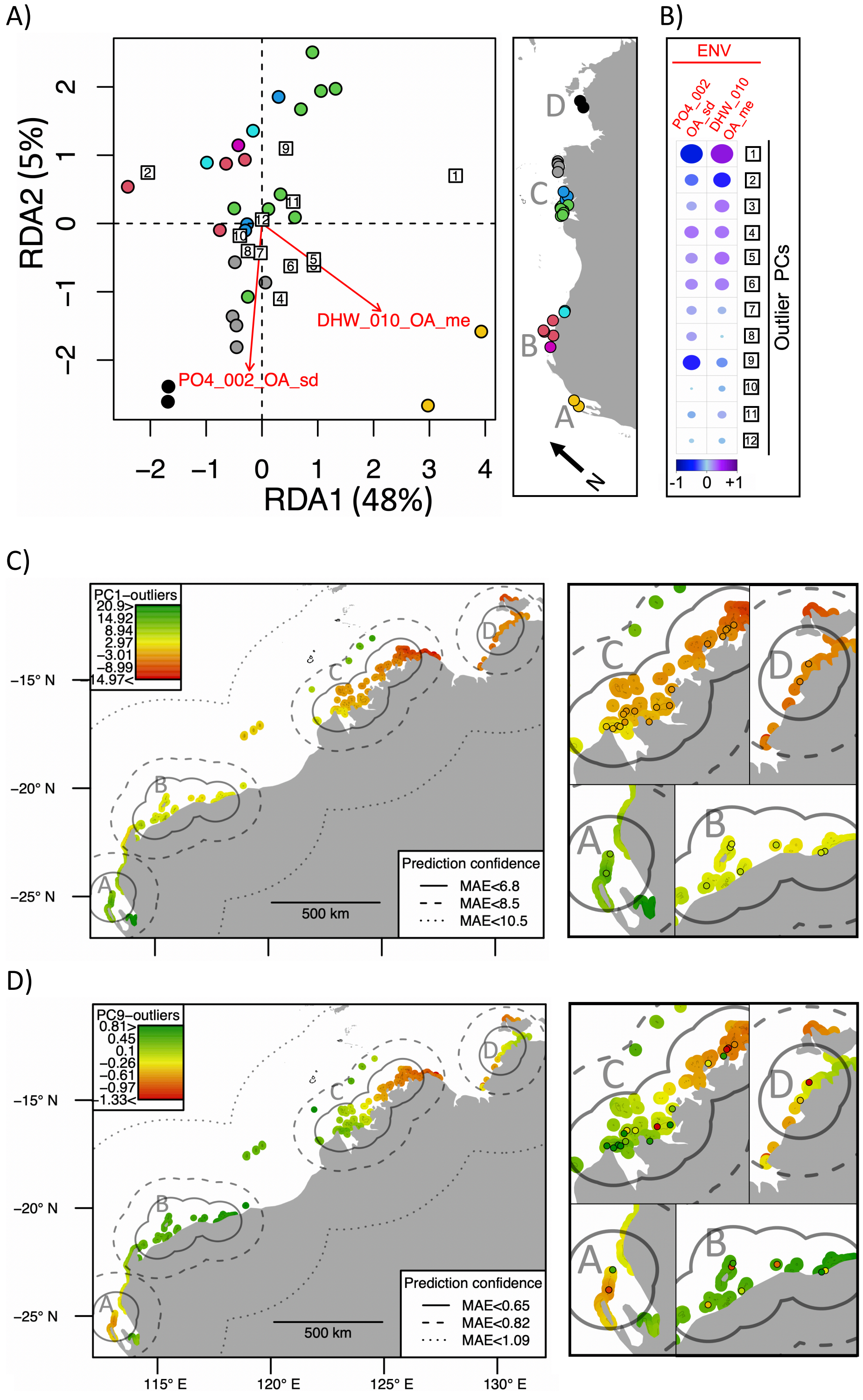

**Supplementary Box 1. Enrichment analysis of biological processes associated with outlier-PCs**

An annotation enrichment analysis was performed to complement the results of the genotype-environment association study. The goal of this analysis was to uncover biological processes putatively controlled by the 62 single nucleotide polymorphism (SNPs) that determine the genetic axes of outlier genetic variation (outlier-PCs, *i.e.* genetic variation not due to neutral structure).

First, we retrieved the nucleotide sequences (on average 57 ± 13 nucleotide per sequence) surrounding each of the 17,007 SNPs characterized through DArT-seq. Next, we ran a similarity search of every sequence against the Uniprot/swissprot protein database (chordata entries, release 2022_01, Boeckmann et al., 2003) using the blastx command from the BLAST command line tool (v. 2.12; Madden & Coulouris, 2008). When a significant match was found (E-value < 0.01), the corresponding SNP was annotated with the gene ontology (GO) terms describing the biological process of the matching protein (Ashburner et al., 2000). In total, 8,423 SNPs were annotated with this method.

We then ran an enrichment analysis of the GO terms annotating the 62 SNPs determining the outlier-PCs using the R package SetRank (v. 1.1; Simillion et al., 2017). For every outlier-PC, we ranked SNPs based on the absolute value of their loading (i.e., higher ranks were attributed to SNPs strongly contributing to the axis of genetic variation). Next, we ran SetRank under the "ranked" mode, to investigate which GO terms were significantly (p-value<0.05; false discoveries-adjusted p-value <0.05) overrepresented among the SNPs attributed with the higher ranks.

The table here below displays the results of the enrichment analysis for the two outlier-PCs most strongly correlated with the environmental variables retained by the RDA: outlier-PC1 (correlated with mean of degree heating week) and outlier-PC9 (correlated with mean degree heating week and standard deviation of phosphate concentration; Supplementary Figure 5). For every outlier-PCs, the table displays the enriched GO term with its description, its occurrence across the annotation of all the SNPs (size) and the p-value of the enrichment test. The complete table for all the outlier-PCs is available in the Supplementary Table 3.

Only one GO term appeared significantly overrepresented among SNPs with a high weight on outlier-PC1: "detection of temperature stimulus". As for outlier-PC9, three terms were found: "inflammatory response", "complement activation, classical pathway" and "innate immune response".

| | *GO term* | *description* | *size* | *p-val* |
|---|---|---|---|---|
| **Outlier-PC1** | GO:0016048 | detection of temperature stimulus | 2 | 0.03 |
| **Outlier-PC9** | GO:0006954 | inflammatory response | 154 | 0.03 |
| | GO:0006958 | complement activation, classical pathway | 27 | 0.03 |
| | GO:0045087 | innate immune response | 180 | 0.03 |

**Supplementary Figure 6. Correlation between environmental variables in case study 2.** The plot displays the levels of average correlation between 32 groups of highly correlated environmental variables (clsXX). The diagonal indicates the average correlation within each group. Below the plot, the variables included in every group of correlated variables are listed. The identifier of the environmental variable involved is shown on top of every plot, and such identifier is defined by (1) the name of the environmental descriptor (CHL: chlorophyll concentration, DHW: degree heating week, FE: iron concentration, NO3: nitrate concentration, O2: oxygen concentration, PH: water pH, PO4: phosphate concertation, SPM: suspended matter concentration, SSS: sea surface salinity, SST: sea surface temperature, CROP: density of cropland areas, LAND: density of surface land, URBA: density of built-up areas, VBD: average boat detection, PDEN: human population density); followed by (2) the buffer size from the survey site, used to calculate the spatial resolution of the variable (002: 2.5 km, 010: 10 km, 025: 25 km and 050: 50 km); followed by (3) the temporal window during which the variable was calculated (DS: dry season, WS: wet season, OA: overall); followed by (4) the statistic used to summarize the environmental variable over time (me: mean, sd: standard deviation).

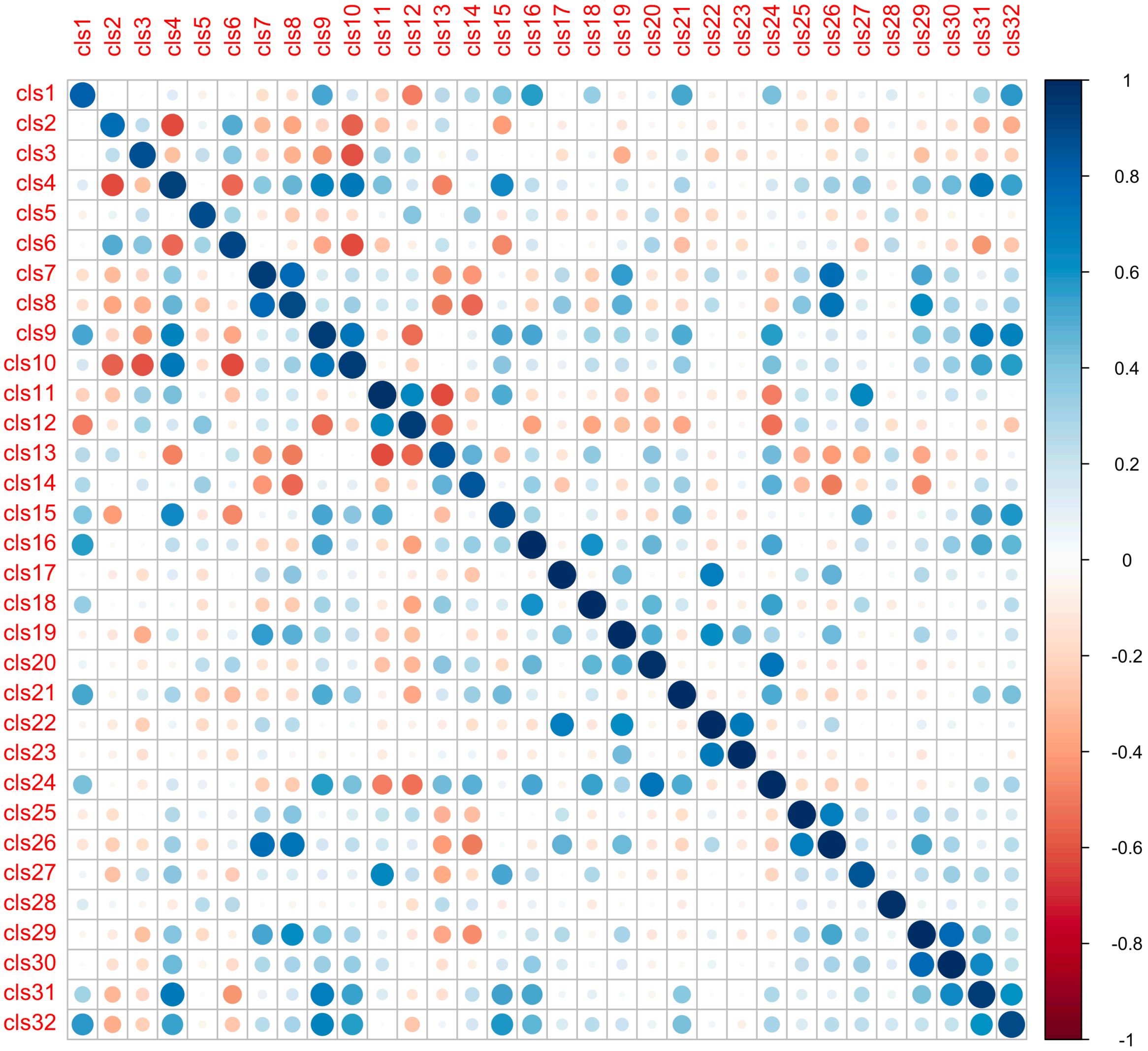


| | |
|---|---|
| **cls1** | CHL_002_OA_me, CHL_002_OA_sd, CHL_002_WS_me, CHL_002_WS_sd, CHL_002_DS_me, CHL_002_DS_sd, CHL_010_OA_me, CHL_010_OA_sd, CHL_010_WS_me, CHL_010_WS_sd, CHL_010_DS_me, CHL_010_DS_sd, CHL_025_OA_me, CHL_025_OA_sd, CHL_025_WS_me, CHL_025_WS_sd, CHL_025_DS_me, CHL_025_DS_sd, CHL_050_OA_me, CHL_050_OA_sd, CHL_050_WS_me, CHL_050_WS_sd, CHL_050_DS_me, CHL_050_DS_sd, SSS_002_OA_me, SSS_002_OA_sd, SSS_002_WS_me, SSS_002_WS_sd, SSS_002_DS_me, SSS_002_DS_sd, SSS_010_OA_me, SSS_010_OA_sd, SSS_010_WS_me, SSS_010_WS_sd, SSS_010_DS_me, SSS_010_DS_sd, SSS_025_OA_me, SSS_025_OA_sd, SSS_025_WS_me, SSS_025_WS_sd, SSS_025_DS_me, SSS_025_DS_sd, SSS_050_OA_me, SSS_050_OA_sd, SSS_050_WS_me, SSS_050_WS_sd, SSS_050_DS_me, SSS_050_DS_sd, SST_002_OA_sd, SST_002_DS_sd, SST_010_OA_sd, SST_010_DS_sd, SST_025_OA_sd, SST_025_DS_sd, SST_050_OA_sd, SST_050_DS_sd |
| **cls2** | DHW_002_OA_me, DHW_002_OA_sd, DHW_002_WS_sd, DHW_002_DS_me, DHW_002_DS_sd, DHW_010_OA_me, DHW_010_OA_sd, DHW_010_WS_sd, DHW_010_DS_me, DHW_010_DS_sd, DHW_025_OA_me, DHW_025_OA_sd, DHW_025_WS_sd, DHW_025_DS_me, DHW_025_DS_sd, DHW_050_OA_me, DHW_050_OA_sd, DHW_050_WS_sd, DHW_050_DS_me, DHW_050_DS_sd, O2_002_OA_me, O2_002_WS_me, O2_002_WS_sd, O2_002_DS_me, O2_010_OA_me, O2_010_WS_me, O2_010_WS_sd, O2_010_DS_me, O2_025_OA_me, O2_025_WS_me, O2_025_WS_sd, O2_025_DS_me, O2_050_OA_me, O2_050_WS_me, O2_050_WS_sd, O2_050_DS_me, PH_002_OA_me, PH_002_WS_me, PH_002_DS_me, PH_002_DS_sd, PH_010_OA_me, PH_010_WS_me, PH_010_DS_me, PH_010_DS_sd, PH_025_OA_me, PH_025_WS_me, PH_025_DS_me, PH_025_DS_sd, PH_050_OA_me, PH_050_WS_me, PH_050_DS_me, PH_050_DS_sd, PO4_002_OA_me, PO4_002_WS_me, PO4_002_DS_me, PO4_010_OA_me, PO4_010_WS_me, PO4_010_DS_me, PO4_025_OA_me, PO4_025_WS_me, PO4_025_DS_me, PO4_050_OA_me, PO4_050_WS_me, PO4_050_DS_me, SST_002_OA_me, SST_002_WS_me, SST_002_WS_sd, SST_002_DS_me, SST_010_OA_me, SST_010_WS_me, SST_010_WS_sd, SST_010_DS_me, SST_025_OA_me, SST_025_WS_me, SST_025_WS_sd, SST_025_DS_me, SST_050_OA_me, SST_050_WS_me, SST_050_WS_sd, SST_050_DS_me |
| **cls3** | DHW_002_WS_me, DHW_010_WS_me, DHW_025_WS_me, DHW_050_WS_me, PO4_002_WS_sd, PO4_010_WS_sd, PO4_025_WS_sd, PO4_050_WS_sd |
| **cls4** | FE_002_OA_me, FE_002_WS_me, FE_002_DS_me, FE_010_OA_me, FE_010_WS_me, FE_010_DS_me, FE_025_OA_me, FE_025_WS_me, FE_025_DS_me, FE_050_OA_me, FE_050_WS_me, FE_050_DS_me |
| **cls5** | FE_002_OA_sd, FE_002_DS_sd, FE_010_OA_sd, FE_010_DS_sd, FE_025_OA_sd, FE_025_DS_sd, FE_050_OA_sd, FE_050_DS_sd |
| **cls6** | FE_002_WS_sd, FE_010_WS_sd, FE_025_WS_sd, FE_050_WS_sd |
| **cls7** | NO3_002_OA_me, NO3_002_OA_sd, NO3_002_WS_me, NO3_002_WS_sd, NO3_002_DS_me, NO3_002_DS_sd, NO3_010_OA_me, NO3_010_OA_sd, NO3_010_WS_me, NO3_010_WS_sd, NO3_010_DS_me, NO3_010_DS_sd, NO3_025_DS_me, NO3_025_DS_sd |

| | |
|---|---|
| **cls8** | NO3_025_OA_me, NO3_025_OA_sd, NO3_025_WS_me, NO3_025_WS_sd, NO3_050_OA_me, NO3_050_OA_sd, NO3_050_WS_me, NO3_050_WS_sd, NO3_050_DS_me, NO3_050_DS_sd |
| **cls9** | O2_002_OA_sd, O2_002_DS_sd, O2_010_OA_sd, O2_010_DS_sd, O2_025_OA_sd, O2_025_DS_sd, O2_050_OA_sd, O2_050_DS_sd |
| **cls10** | PH_002_OA_sd, PH_010_OA_sd, PH_025_OA_sd, PH_050_OA_sd |
| **cls11** | PH_002_WS_sd, PH_010_WS_sd, PH_025_WS_sd, PH_050_WS_sd |
| **cls12** | PO4_002_OA_sd, PO4_002_DS_sd, PO4_010_OA_sd, PO4_010_DS_sd, PO4_025_OA_sd, PO4_025_DS_sd, PO4_050_OA_sd, PO4_050_DS_sd |
| **cls13** | SCV_002_OA_me, SCV_002_OA_sd, SCV_002_WS_me, SCV_002_DS_me, SCV_002_DS_sd, SCV_010_OA_me, SCV_010_OA_sd, SCV_010_WS_me, SCV_010_DS_me, SCV_010_DS_sd, SCV_025_OA_me, SCV_025_OA_sd, SCV_025_WS_me, SCV_025_DS_me, SCV_025_DS_sd, SCV_050_OA_me, SCV_050_OA_sd, SCV_050_WS_me, SCV_050_DS_me, SCV_050_DS_sd |
| **cls14** | SCV_002_WS_sd, SCV_010_WS_sd, SCV_025_WS_sd, SCV_050_WS_sd |
| **cls15** | SPM_002_OA_me, SPM_002_OA_sd, SPM_002_WS_me, SPM_002_WS_sd, SPM_002_DS_me, SPM_002_DS_sd, SPM_010_OA_me, SPM_010_OA_sd, SPM_010_WS_me, SPM_010_WS_sd, SPM_010_DS_me, SPM_010_DS_sd, SPM_025_OA_me, SPM_025_OA_sd, SPM_025_WS_me, SPM_025_WS_sd, SPM_025_DS_me, SPM_025_DS_sd, SPM_050_OA_me, SPM_050_OA_sd, SPM_050_WS_me, SPM_050_WS_sd, SPM_050_DS_me, SPM_050_DS_sd |
| **cls16** | PDEN_002_me |
| **cls17** | PDEN_010_me |
| **cls18** | PDEN_010_sd |
| **cls19** | PDEN_025_me |
| **cls20** | PDEN_025_sd, PDEN_050_me, URBA_025_me, URBA_050_me, URBA_050_sd, VBD_010_me, VBD_010_sd, VBD_025_me, VBD_025_sd |
| **cls21** | PDEN_050_sd |
| **cls22** | URBA_010_me |
| **cls23** | URBA_010_sd |
| **cls24** | URBA_025_sd, VBD_050_me, VBD_050_sd |
| **cls25** | CROP_002_me |
| **cls26** | CROP_002_sd |
| **cls27** | CROP_010_me, CROP_010_sd, CROP_025_me, CROP_025_sd, CROP_050_me, CROP_050_sd |
| **cls28** | VBD_002_me, VBD_002_sd |
| **cls29** | LAND_002 |
| **cls30** | LAND_010 |
| **cls31** | LAND_025, LAND_050 |
| **cls32** | BATHY_002, BATHY_010, BATHY_025, BATHY_050 |

**Supplementary Figure 7. Absolute error from spatially-explicit cross validations of case study 2.** The boxplot displays the distribution of the absolute error of the outlier-PC1 (A) and outlier-PC9 (B) predicted by the cross-validation procedure. Each box corresponds to the iterations performed under a given value of the threshold distance D, which was used to group together survey sites that are spatially close in the spatially-explicit cross validation procedure.

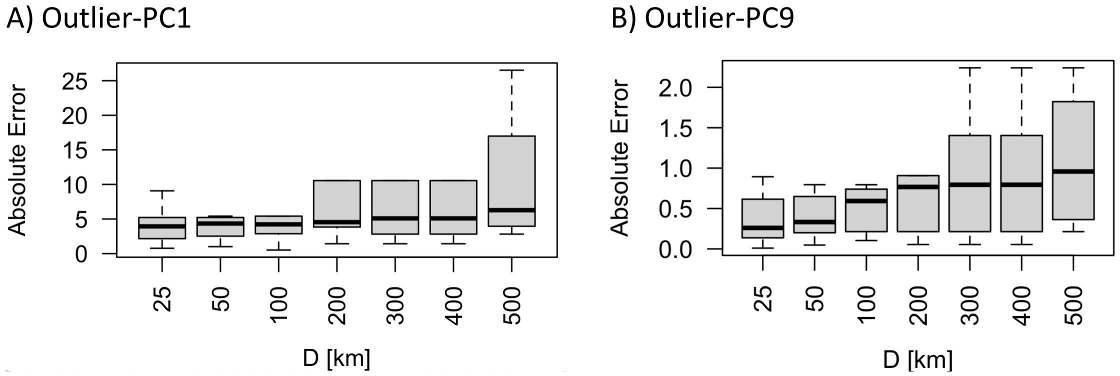
